# Connectivity analysis of single cell RNA-sequencing derived transcriptional signature of lymphangioleiomyomatosis

**DOI:** 10.1101/2020.09.30.320473

**Authors:** Naim Al Mahi, Erik Y. Zhang, Susan Sherman, Jane J. Yu, Mario Medvedovic

**Affiliations:** Division of Biostatistics and Bioinformatics, Department of Environmental and Public Health Sciences, University of Cincinnati College of Medicine, Cincinnati, OH 45267, USA; Department of Biomedical Informatics, University of Cincinnati College of Medicine, Cincinnati, OH 45267, USA; Division of Pulmonary, Critical Care and Sleep Medicine, University of Cincinnati College of Medicine, Cincinnati, OH 45267, USA; The LAM Foundation, Cincinnati, OH 45242, USA

## Abstract

Lymphangioleiomyomatosis (LAM) is a rare pulmonary disease affecting women of childbearing age that is characterized by the aberrant proliferation of smooth-muscle (SM)-like cells and emphysema-like lung remodeling. In LAM, mutations in TSC1 or TSC2 genes results in the activation of the mechanistic target of rapamycin complex 1 (mTORC1) and thus sirolimus, an mTORC1 inhibitor, has been approved by FDA to treat LAM patients. Sirolimus stabilizes lung function and improves symptoms. However, the disease recurs with discontinuation of the drug, potentially because of the sirolimus-induced refractoriness of the LAM cells. Therefore, there is a critical need to identify remission inducing cytocidal treatments for LAM. Recently released Library of Integrated Network-based Cellular Signatures (LINCS) L1000 transcriptional signatures of chemical perturbations has opened new avenues to study cellular responses to existing drugs and new bioactive compounds. Connecting transcriptional signature of a disease to these chemical perturbation signatures to identify bioactive chemicals that can “revert” the disease signatures can lead to novel drug discovery. We developed methods for constructing disease transcriptional signatures and performing connectivity analysis using single cell RNA-seq data. The methods were applied in the analysis of scRNA-seq data of naïve and sirolimus-treated LAM cells. The single cell connectivity analyses implicated mTORC1 inhibitors as capable of reverting the LAM transcriptional signatures while the corresponding standard bulk analysis did not. This indicates the importance of using single cell analysis in constructing disease signatures. The analysis also implicated other classes of drugs, CDK, MEK/MAPK and EGFR/JAK inhibitors, as potential therapeutic agents for LAM.

## Introduction

Lymphangioleiomyomatosis (LAM) is a progressive interstitial lung disease predominantly affects young females of reproductive age carrying an inherited disorder called Tuberous Sclerosis Complex (TSC) or due to a sporadic form without any evidence of a genetic disease^1–3^. This uncommon complex disease is caused by activation of the mechanistic target of rapamycin complex 1 (mTORC1) through inactivating mutations in tumor suppressor genes TSC1 or TSC2 which is directly associated with unrestricted cell growth^3,4^. Besides smooth muscle (SM) cell proliferation and emphysema-like lung remodeling^5^, LAM also results from the infiltration of neoplastic cells containing both SM and melanocyte lineage cells^6,7^ leading to interstitial cystic lung destruction^8^.

Currently, mTORC1 inhibitor sirolimus is the only drug approved by the Food and Drug Administration (FDA) which improves pulmonary dysfunction and decelerates LAM progression in most patients^9^. However, sirolimus treatment does not lead to progression free survival and has a cytostatic rather than a cytocidal effect. Lung function decline resumes following drug discontinuation and thus uninterrupted drug exposure is required for prolonged benefit^9,10^. The drug cannot completely eliminate LAM cells potentially because chronic exposure to sirolimus induces refractoriness and resistant behavior of the mTORC1-hyperactive LAM cells^11^. Therefore, it is urgent to identify remission-inducing and durably effective therapeutic agents for LAM.

As an alternative to *de novo* drug discovery, identifying new therapeutic uses of the existing drugs by leveraging large compendia of biomedical data, also known as drug repositioning, has been used as a potential tool in drug discovery and development^12–14^. In the connectivity map (CMap) drug repositioning^15^, transcriptional signature of disease is constructed by differential gene expression analysis between the diseased tissue or cells and the control. The negative correlation between the transcriptional disease signature and the transcriptional signature of the drug treatment is used to identify drugs capable of “reversing” the disease process to be used as potential therapeutics. For example, histone deacetylase (HDAC) inhibitor vorinostat, which is known to treat cutaneous T-cell lymphoma, has been shown to be effective in treating gastric cancer^16^ or drug topiramate has been identified as a potential candidate to treat inflammatory bowel disease (IBD) by comparing gene expression signatures of IBD against drug perturbational signatures^17^. The most recent edition of the connectivity map library, generated by the integrated network-based cellular signatures (LINCS) project, catalogues transcriptional signatures of more than 20,000 drugs and uncharacterized small chemicals across 77 cell lines facilitating drug repositioning and identification of new therapeutic agents^18,19^.

With the recent progress of next generation sequencing technologies, single-cell RNA-seq (scRNA-seq) has emerged as a powerful tool to investigate inter-cellular heterogeneity at single cell level. The gene expression dynamics of individual cells provides means to study complex disease mechanisms at an unprecedented resolution. Although considerable research has been devoted to using bulk transcriptional signatures for computational drug repositioning, methodologies for connecting diseases, genes, and drugs using scRNA-seq data are lacking. In this paper we develop the complete protocol for performing connectivity analysis using scRNA-seq data, including signatures construction and connectivity analysis with individual drug signatures as well as the whole classes of drugs with the same mechanism of action. We use the new methods to perform connectivity analysis of LAM scRNA-seq signatures. Our analyses confirm therapeutic effect of currently used drugs and provides additional drug candidates. Importantly, we demonstrate that these results are contingent on use of scRNA-seq data and our methods for constructing single cell disease signature and would not be possible by connectivity analysis of standard bulk RNA-seq disease signatures.

## Results

### Overview of scRNA-seq connectivity analysis

Conventional transcriptome profiling methods such as bulk RNA-seq relies on averaging molecular signals across a large population of cells. This can lead to missing key expression features of a small subpopulation of cells that may be crucial for disease progression and response to target therapies. The goal of our analysis is to construct a transcriptional signature of disease-critical cells which may represent a small fraction of profiled cells. Our analysis identifies the disease-critical cell subpopulations and constructs the disease signature by comparing the expression profile of disease-critical cells to the matched cell type in the control non-diseased tissue. Our central hypothesis is that such a single cell disease signature will factor out the cell-type to cell-type differences, and will facilitate identification of effective therapeutics when the standard connectivity analysis of bulk disease signatures fails.

The analytical workflow of scRNA-seq signature construction and connectivity analysis proceeds as: (1) Cluster analysis of disease and controls samples; (2) Construct cluster annotating signature (CAS) for each cluster in the disease sample and identification of the disease-critical cell subpopulation using the panel of disease marker genes; (3) Identify matching control cell populations in the non-diseased sample; (4) Construct disease characterizing signature (DCS) by comparing the disease-critical cells with the matched control cells; (4) “Connect” DCS to LINCS-L1000 chemical perturbational signatures. Details of each step are provided in the Methods, outlined in Supplementary Figure 1, and illustrated through analysis of LAM samples.

### Signature construction and connectivity analysis of naïve LAM

scRNA-seq data were generated using 10x Chromium platform on dissociated lungs from one naïve LAM patient (LAM1), one sirolimus treated LAM patient (LAM2), and one normal patient (WT) respectively, and has been previously described and analyzed^20^. In total, 19,384 cells (7,244 cells from LAM1, 6,545 cells from LAM2, and 5,595 cells from WT) were included in the downstream analyses after filtering out low quality cells from each sample separately (Methods), with an average number of detected genes (UMI>0) of 2,089, 2,466, and 1,564 per cell in LAM1, LAM2, and WT respectively (Supplementary Figure 2). The analytical workflow outlined above were carried out for LAM1 and LAM2 samples separately.

### Cluster analysis of naïve LAM and wild-type samples

Single-cell clustering was performed for naïve LAM (LAM1) and wild-type (WT) sample individually. We employed graph-based clustering implemented in Seurat3^21^, which identified 19 clusters in each of the samples that are visualized using t-Distributed Stochastic Neighbor Embedding (t-SNE) plots (Methods; **Error! Reference source not found.**A).

### Construction of cluster annotating signatures

To construct cluster annotating signatures (CAS), pairwise comparisons for each cluster was conducted and then combined into a single cluster specific signature (Methods; Supplementary Figure 1). The top most significantly (False discovery rate (FDR) <0.05) up-regulated genes, namely cluster annotating signature (CAS) were then used to annotate cell clusters by cell-types or tissues. This step was iterated for each cluster separately.

To identify disease-critical cell sub-population, we utilized a set of 8 marker genes identified as the markers of LAM from the literature (Figure 1**Error! Reference source not found.**B; Supplementary Table 1). All the markers were exclusively highly expressed in cluster 16 of LAM1 (Figure 1B), and this cluster was the only one whose signature was enriched for expression of the marker genes (Figure 1C; Supplementary Table 2) indicating that the cluster (herein denoted as LAM1_cluster16_) consists of LAM cells.

**Figure 1.**
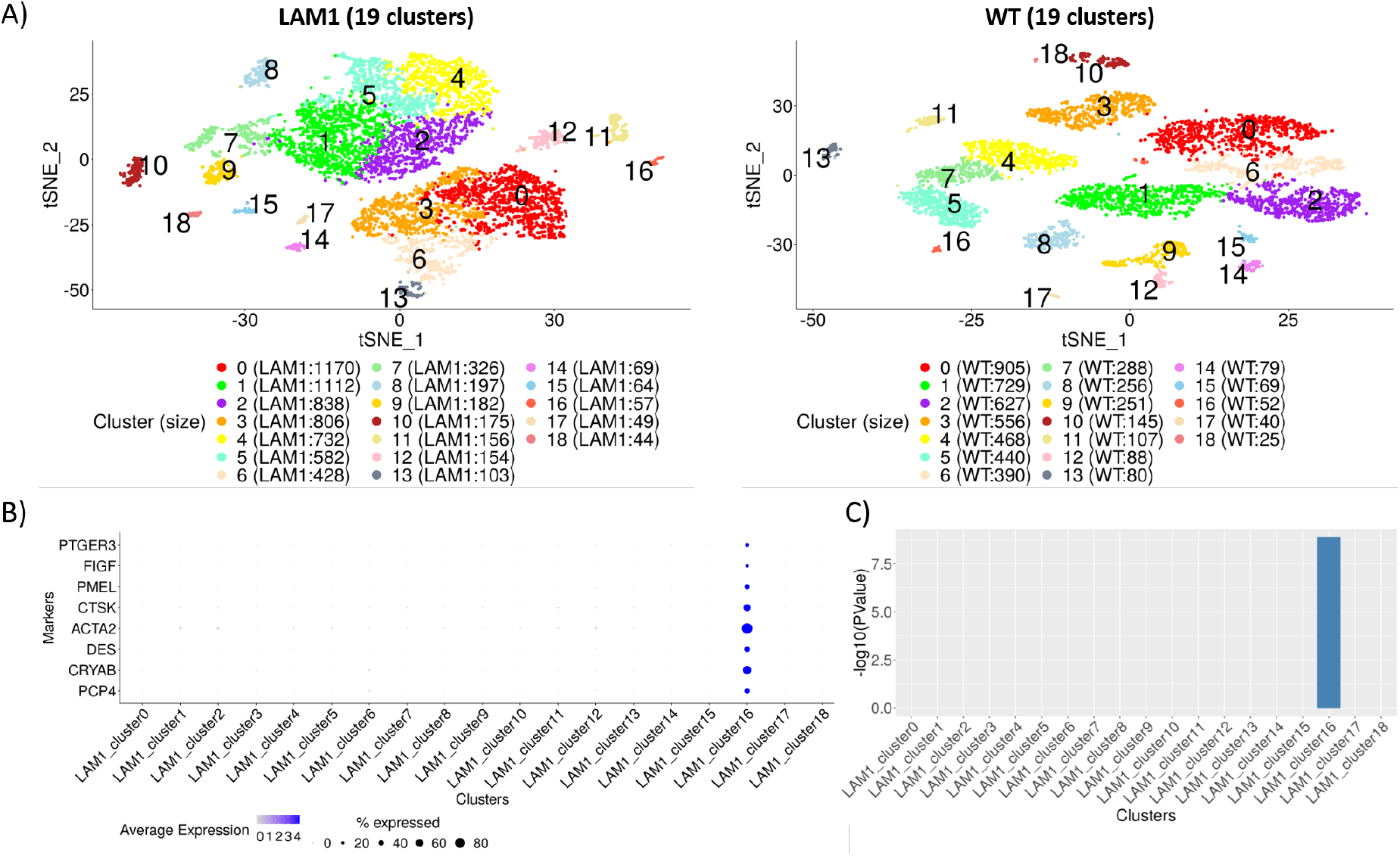
Cluster analysis of scRNA-seq samples. **A)** Unsupervised clustering of 7,244 cells from LAM1 (top panel) and 5,595 cells from wild-type (WT) sample (bottom panel) are represented in twodimensional t-SNE plots with perplexity 30. A total of 19 clusters were identified in each sample using Seurat’s graph-based clustering initialized with top principal components with largest variances. Clusters are colored and labeled distinctively and the number of cells in each cluster is noted inside the parenthesis in the legends. **B)** Expression of known LAM markers was used to identify the cluster of LAM cells, with the size of the dot representing the percentage of cells expressed and color is proportional to the average expression of the genes. All the 8 markers show moderate to high expression in at least 30% cells in cluster 16 of LAM1. **C)** Marker enrichment was conducted using Fisher’s exact test based on the significantly (FDR<0.05) differentially expressed (DE) genes from each of the cluster annotating signatures of LAM1. All the markers were significantly DE only in cluster 16, whereas none of the markers were significant in any other cluster.

To further characterize cells in different clusters, we performed enrichment analysis of the top 200 most significantly up-regulated genes from each cluster for cell type marker from three databases: Human cell landscape (HCL)^22^, cellMarker (CM)^23^, and PanglaoDB (PDB)^24^, and the tissue markers derived from the gene atlas dataset^25^. Top 3 most significantly (FDR<0.05) enriched tissue and cell-type categories with log odds ratio above 1.5 from each cluster were selected for each cluster and associations between the clusters and cell and tissue type are summarized in the Supplementary Figure 3. The analysis implicated clusters of different kinds of epithelial, endothelial, and immune cells. The cells implicated by the CAS of LAM1_cluster16_ cells was enriched for markers of mesenchymal cells and uterus, uterus-corpus and appendix tissue signatures (Supplementary Figure 3).

### Construction of disease characterizing signature

Disease characterizing signature of LAM1 was constructed by comparing LAM1_cluster16_ with the transcriptionally analogous WT clusters. Comparing LAM1_cluster16_, which is a cluster of smooth-muscle like cells, to any WT cluster such as, B cells, T cells, or endothelial cells, would increase noise and might not show the signal pertinent to transcriptional changes in LAM cells. Therefore, selecting only the WT clusters that were most similar to LAM1_cluster16_ detected relevant transcriptional changes in LAM cells compared to the equivalent non-diseased cells.

The analysis of overlaps between the LAM1_cluster16_ CAS and CASes of all WT clusters identified cells in WT clusters 9 and 12 (Figure 2A; Figure 2B) as being the most similar to the LAM cells in LAM1_cluster16_. Single cell disease characterizing signature (DCS) of LAM was then constructed by differential gene expression analysis between cells in LAM1_cluster16_ and cells in WT clusters 9 and 12. To illustrate the advantages of the single cell DCS, we also constructed pseudo-bulk signature of LAM1 by differential expression between all LAM1 cells and all WT cells (Methods). This signature mimics the signature that would be obtained by the bulk RNA-seq analysis.

**Figure 2.**
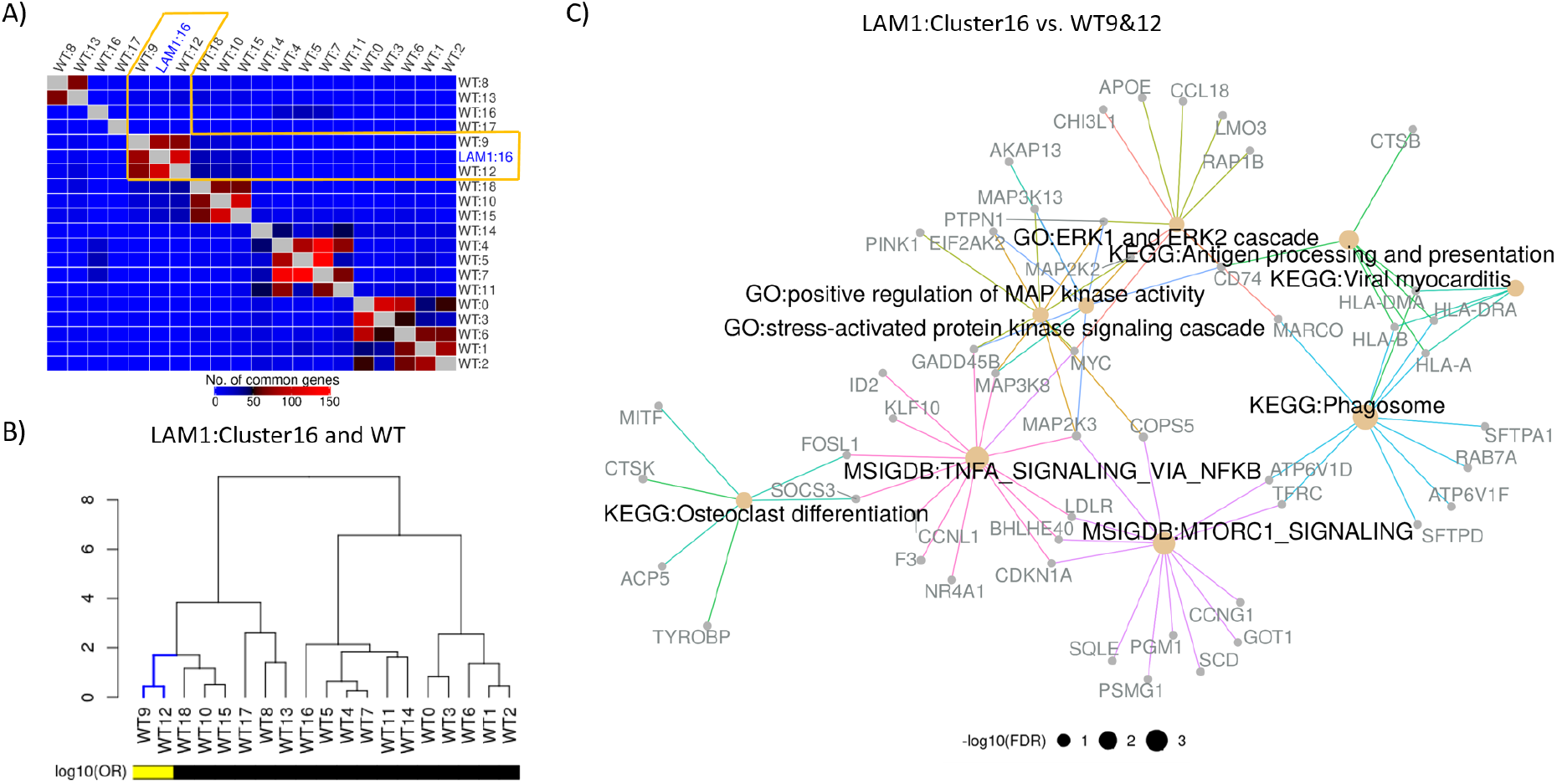
Construction and functional enrichment of disease characterizing signature. **A)** WT clusters were matched with the LAM cluster in terms of top 200 most significantly (FDR<0.05) up-regulated genes from each of the CAS. Unsupervised hierarchical clustering revealed sub-clusters of LAM and WT clusters, where LAM1_cluster16_ was clustered with WT clusters 9 and 12. **B)** Significance of the overlaps between LAM and WT cell clusters based on the significantly (FDR<0.05) up-regulated genes were assessed via Fisher’s exact test. Cluster similarity were measured using log10 odds ratio and hierarchical clustering of LAM1_cluster16_ vs. WT is visualized via dendrograms. Log_10_ odds ratio (OR) of 1 or more is indicated by the yellow color. **C)** Disease characterizing signatures of LAM were constructed by comparing LAM1_cluster16_ with the WT cluster 9 and 12. Functional enrichment of top 200 most significantly (FDR<0.05) up-regulated genes was carried out in terms of KEGG/MSigDB (Hallmark)/GO (Biological processes) categories. Selected functional classes based on the cutoff of FDR adjusted P-values<0.1 are represented by different edge colors and size of the node is proportional to negative logarithm of FDR adjusted P-value.

The pathway analysis of the LAM single cell DCS against GO^26^, KEGG^27^, and MSigDB (Hallmark)^28^ gene sets via clusterProfiler^29^, implicated MTORC1 signaling hallmark gene sets as being enriched in the DCS (Figure C), along with gene sets pathways associated with cell proliferation, invasion, and metastasis. Although, most of these signaling pathways are known features of LAM, identifying their activity within the LAM cell populations based on a transcriptional signature is not a trivial task. The analysis of the pseudo-bulk LAM signature does not reveal increased MTOR signaling (Supplementary Figure 4A), demonstrating the increasing precision of our DCS in comparisons to a typical signature constructed from bulk tissue profiling.

### Connectivity analysis

We developed a protocol to perform the connectivity analysis of a DCS against 143,374 LINCS signatures (Methods) in response to treatment with 15,349 chemical perturbagens (CP), and identify potential drug or small molecule candidates for treatment of LAM. We developed an analytical framework to connect LAM signature to LINCS CP signatures and identify MOA of the drugs/small molecules with connected signatures (Methods). Briefly, single cell DCS is correlated with individual LINCS CP signatures (Figure **3**A). The enrichment of signature with high negative correlations among CPs with a specific MOA was assessed using small-sample bias corrected logistic regression. We identified several cell proliferation and pro-survival pathway targets in LAM1. Most enriched MOA categories included both MTOR inhibitors, dual inhibition of PI3K/MTOR, and CDK inhibitors (Figure **3**B).

**Figure 3.**
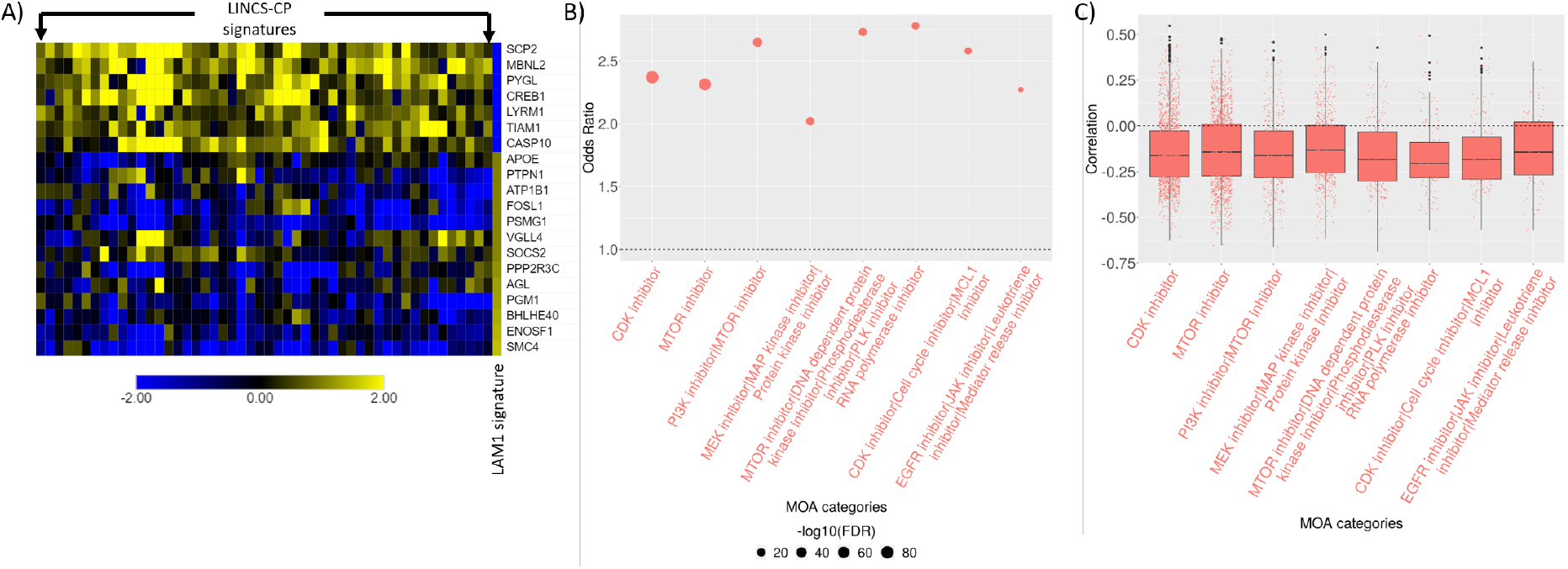
Connectivity analysis of naïve LAM signature. **A)** Top 250 most up/down regulated genes from LAM characterizing signature was selected and matched with 978 LINCS-L1000 landmark genes. Pearson’s correlation were computed between each of the LINCS-CP and LAM signature. Top 50 LINCS-CP signatures most negatively correlated with the LAM signature (columns) with the corresponding matched genes (rows) are presented via heatmap. **B)** Odds ratios of the top most enriched MOA categories are shown via dot plot where the size of the dots represent the significance of the MOA categories with a bigger dot indicating lower FDR adjusted P-value. MOA categories were selected based on odds ratio>2, –log10(FDR)>7, and at least 100 signatures in any MOA category. **C)** Distribution of the overall signature correlations associated with each of the MOA categories are demonstrated via box-and-whisker plots. Each dot represents a LINCS-CP signature and negative correlations indicate the potential of the drug mechanisms to revert the LAM signature.

Given the known etiology of LAM, and the use of the sirolimus MTOR inhibitor in the treatment of LAM, ability of MTOR inhibitors to reverse the LAM is expected and also in line with the functional analysis results from the previous section. However, the same connectivity analysis repeated on the pseudo-bulk LAM signature fails to identify MTOR inhibitors as putative therapeutics (Supplementary Table 5). This again demonstrates the importance of the carefully constructed single cell DCS for the successful connectivity analysis. We found sirolimus, AZD-8055, OSI-027, and WYE-125132 showing consistently strong negative correlation across all the dosages with LAM1 DCS (Supplementary Figure 5A).

Cyclin-dependent kinase inhibitors (CKI) play a vital role in controlling cell cycle progression and cell proliferation by inhibiting specific cyclin/cyclin-dependent kinase complexes^30,31^. CDK1/2 inhibitors CGP-60474, PHA-793887, and alvocidib and CDK4/6 inhibitor palbociclib shows strong negative correlation with LAM1 single cell DCS across different concentrations and cell lines (Supplementary Figure 5A). Functional enrichment of the LAM DCS identified biological processes and pathways related to MAP kinase signaling (Figure **2**C) which was also supported by our connectivity analysis with MEK/MAP kinase/protein kinase inhibitors being implicated as putative therapeutic agents. Estrogen-induced activation of MAPK signaling is associated with enhanced cell proliferation^32^ and survival of LAM cells^33^. Estrogen-increased the expression of oncogene c-MYC, which plays a critical role in cell cycle progression by suppressing p21^Cip1^ expression^34^, in LAM cells (Figure **2**C) and might induce MAPK signal transduction pathways^32,35^. Moreover, inhibition of mTORC1 is known to activate MAPK signaling cascade^36^ which may implicate that combined inhibition of mTORC1 and MAPK can serve as an alternative treatment strategy possibly with better prognosis than sirolimus based monotherapy^37^. Furthermore, signatures from breast cancer cell lines were strongly negatively correlated with LAM1 DCS (Supplementary Figure 5B). Several other pathway inhibitors related to cell proliferation and survival such as HSP, EGFR/JAK, AKT, VEGFR, IGF-1, and HDAC were also associated with LAM1 DCS (Supplementary Table 3).

### Signature construction and connectivity analysis of sirolimus treated LAM

Similar to naïve LAM, we repeated the analytical workflow for sirolimus treated LAM sample (LAM2). The clustering algorithm identified 19 clusters in LAM2 (Figure 4A) and we used LAM marker genes to identify LAM cells in LAM2. However, unlike LAM1, the expression of LAM markers was not localized in any particular cluster, and cells expression were dispersed in all clusters (Figure 4A) making it impossible to identify a single LAM cluster. Marker enrichment in LAM2 cluster further showed no statistical significance in any LAM2 cluster (Supplementary Table 2). As an alternative strategy, we integrated LAM1 and LAM2 cells and re-clustered them. A total of 13,789 cells from LAM1 and LAM2 were combined using Seurat’s^21^ implementation of multiple dataset integration and 18 clusters were detected (Figure 4B). Majority of the markers were comparatively highly expressed in both LAM1 and LAM2 part of cluster 16 (Figure 4B) which was further supported by the enrichment of LAM markers in the joint clusters (Supplementary Table 2). Contractile proteins such as, α-smooth-muscle actin (ACTA2) and desmin (DES), protease cathepsin-K (CTSK), melanocyte protein (PMEL), and alpha-crystallin B chain (CRYAB) were highly expressed in majority of the cells in joint cluster 16. All the 57 cells from LAM1_cluster16_ were also present in the joint cluster 16. The 12 LAM2 cells in the joint cluster 16 were assumed to represent LAM cells in the LAM2 samples and were denoted as LAM2_joint-cluster16_.

**Figure 4.**
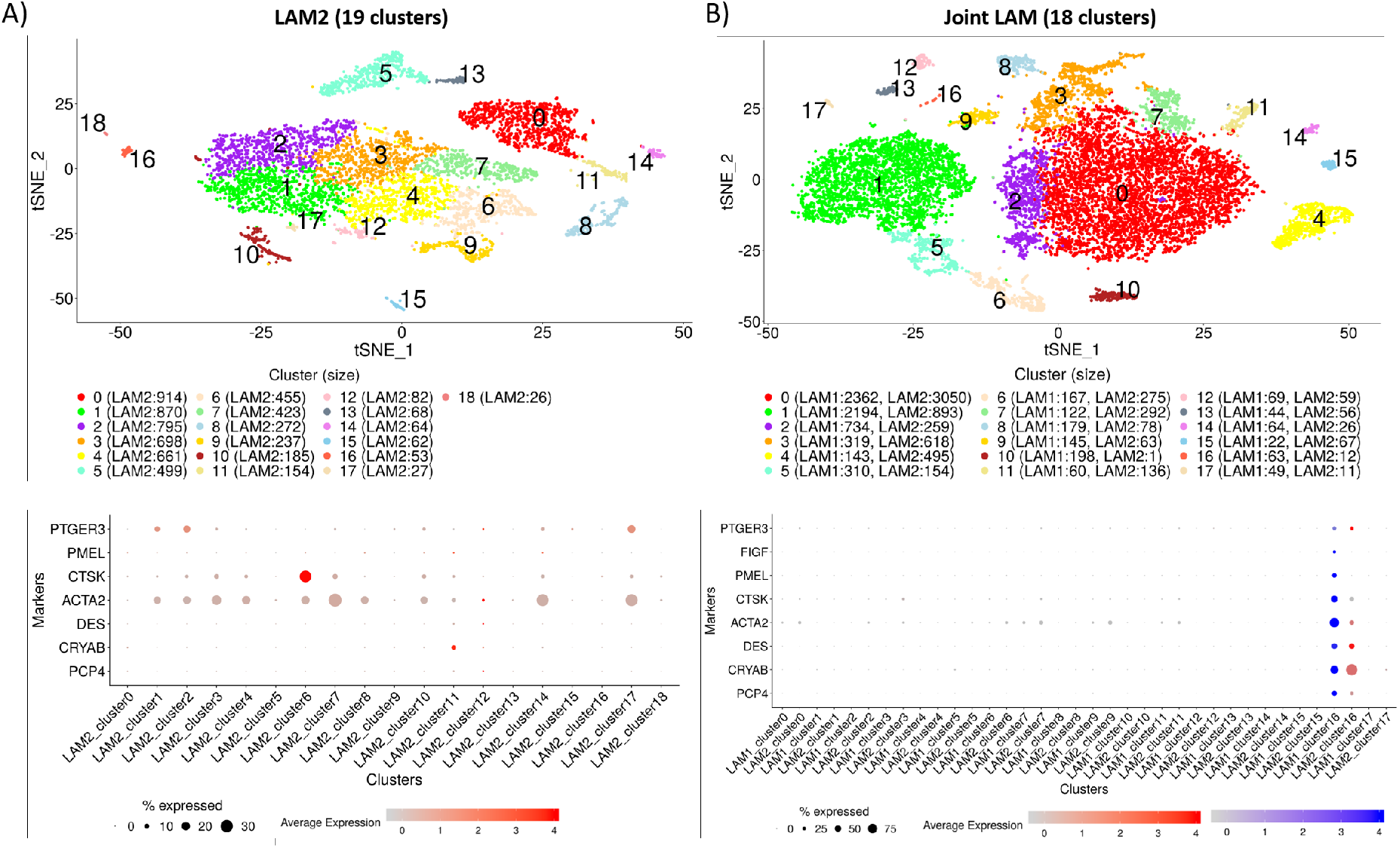
Cluster analysis of LAM2. **A)** Unsupervised clustering of 6,545 cells from LAM2 are represented in two-dimensional t-SNE plots (top panel) with perplexity 30. A total of 19 clusters were identified in each sample using Seurat’s graph-based clustering initialized with top principal components with largest variances. Expression of known LAM markers was used to identify the cluster of LAM cells (bottom panel), with the size of the dot representing the percentage of cells expressed and color is proportional to the average expression of the genes. **B)** Integrated clustering of 13,789 cells from both LAM1 (7,244 cells from LAM1) and LAM2 (6,545 cells from LAM2) identified 18 clusters where each cluster consists of both LAM1 and LAM2 cells (top panel). Seurat’s implementation of integrated clustering was used to identify common cell clusters between LAM1 and LAM2. Clusters are colored and labeled uniquely and the number of cells in each cluster is noted inside the parenthesis in the legends. 6 out of 8 LAM markers show moderate to high expression in at least 25% cells in both LAM1 (63 cells) and LAM2 (12 cells) of cluster 16 (bottom panel).

Cluster annotating signatures of the joint clusters showed similar cell and tissue types as in LAM1 analysis (Supplementary Figure 6). Cluster annotating signatures were further used to find the WT clusters akin to LAM2_joint-cluster16_. Similar to LAM1_cluster16_, WT cluster 9 and 12 had maximum number of overlapping genes with LAM2_joint-cluster16_ (Figure 5A; Figure 5B). The single cell DCS of LAM2 cells was constructed by differential gene expression analysis between cells in LAM2_joint-cluster16_ and the WT clusters 9 and 12. The pathway analysis of the LAM2 DCS identified gene sets associated with the regulation of cell-cell adhesion, response to interferon gamma and tumor necrosis factor, but not MTOR signaling (Figure 5C).

**Figure 5.**
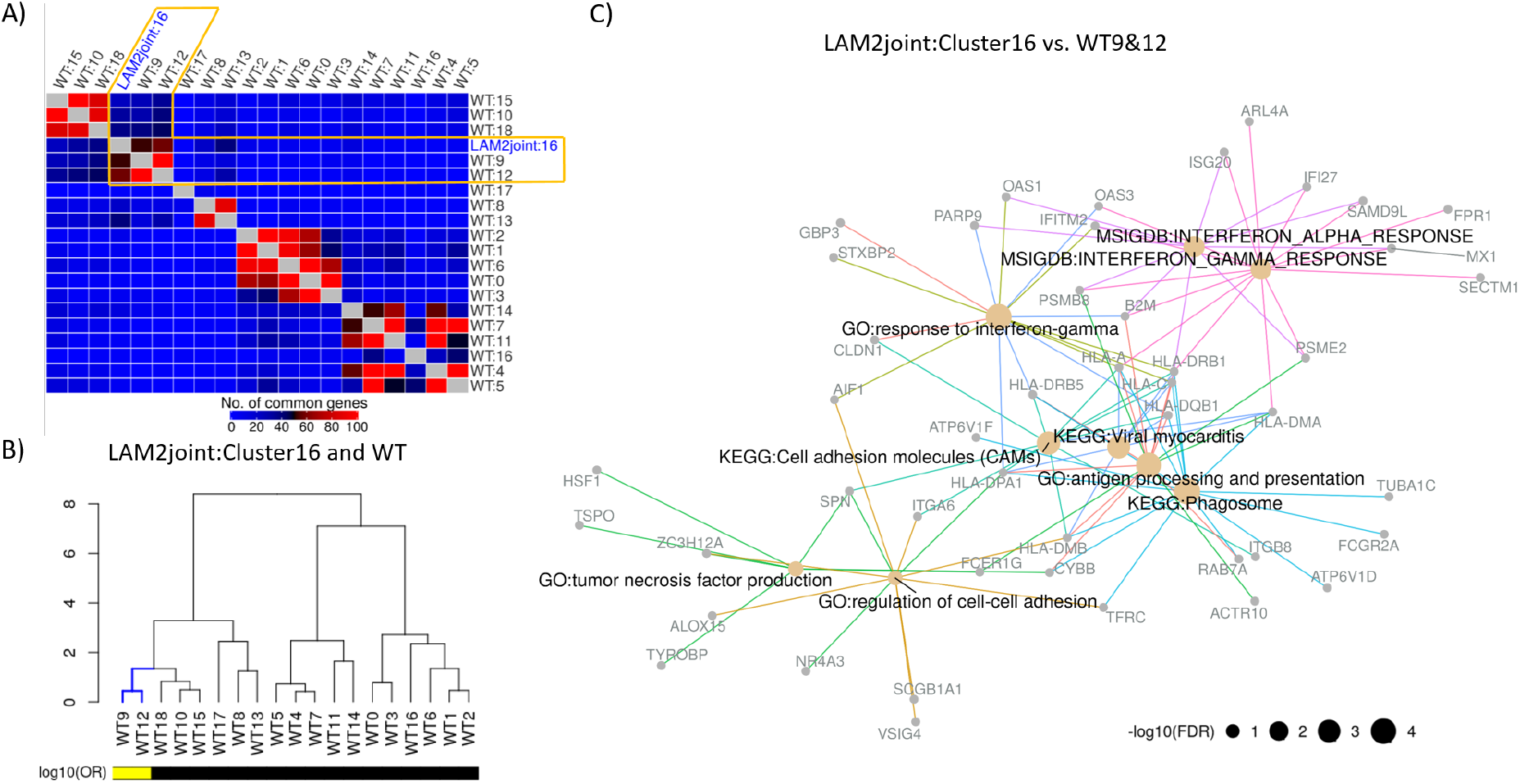
Construction and functional enrichment of disease characterizing signature from LAM2. **A)** WT clusters were matched with the LAM2_joint-cluster16_ cluster in terms of top 200 most significantly (FDR<0.05) up-regulated genes from each of the CAS. Unsupervised hierarchical clustering revealed sub-clusters of LAM and WT clusters, where LAM2_joint-cluster16_ was clustered with WT clusters 9 and 12. **B)** Significance of the overlaps between LAM and WT cell clusters based on the significantly (FDR<0.05) up-regulated genes were assessed via Fisher’s exact test. Cluster similarity were measured using log_10_ odds ratio and hierarchical clustering of LAM1_cluster16_ vs. WT is visualized via dendrograms. Log_10_odds ratio (OR) of 1 or more is indicated by the yellow color. **C)** Disease characterizing signatures of LAM were constructed by comparing LAM2_joint-cluster16_ with the WT cluster 9 and 12. Functional enrichment of top 200 most significantly (FDR<0.05) up-regulated genes was carried out in terms of KEGG/MSigDB (Hallmark)/GO (Biological processes) categories. Selected functional classes based on the cutoff of FDR adjusted P-values<0.05 and odds ratio>2 are represented by different edge colors and size of the node is proportional to negative logarithm of FDR adjusted P-value.

Connectivity analysis of LAM2 DCS (Figure 6A) revealed several MOA categories including single-agent proteasome inhibitors, dual inhibition of NF-κB pathway/proteasome inhibitors and HSP inhibitors. (Figure 6B). Mutation of TSC2 and its leading activation of MTORC1 upregulates the proteasome^38^ which may facilitate estrogen enhanced survival of tumor cells^39,40^. MTOR also activates NF-κB^41^, a major regulator of cell survival, pro-inflammatory cytokines such as TNF-α, and cell adhesion molecules which may allow LAM cells to survive^4,42^. We also found response to interferon gamma and cell adhesion molecules in the functional enrichment of LAM2 DCS (Figure 5C) which might activate NF-κB and supports the anti-apoptotic behavior of the LAM cells. Proteasome inhibitor, which inhibits NF-κB activation, has been found to reduce estrogen mediated survival of TSC2-null cells in LAM^40^ and was one of the top hits in our connectivity analysis with LAM2 DCS. Signatures of tyrosine kinase and cyclooxygenase inhibitor drugs were also implicated (Figure 6B and Figure 6C). Interestingly, several drugs related to this MOA, such as multi-kinase inhibitor imatinib, Src inhibitor Saracatinib, and Cyclooxygenase inhibitor Celecoxib are being currently tested in clinical trials as LAM therapeutics confirming the relevance of the connectivity analysis results. We also found MTOR inhibitors as one of the top enriched MOA categories although with relatively low strength of association (odds ratios) (Figure 6B).

**Figure 6.**
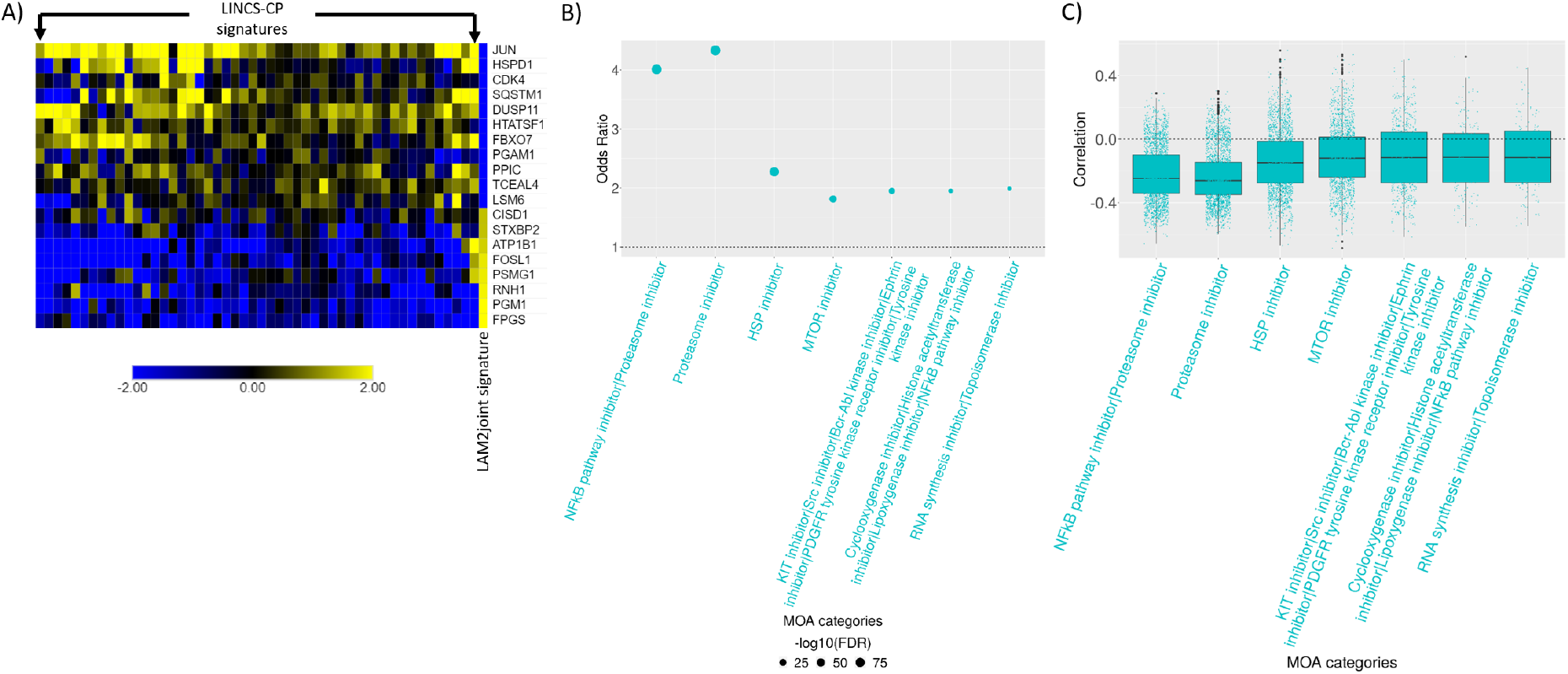
Connectivity analysis of sirolimus treated LAM signature. **A)** Top 250 most up/down regulated genes from LAM2 DCS signature was selected and matched with 978 LINCS-L1000 landmark genes. Pearson’s correlation was computed between each of the LINCS-CP and LAM signature. Top 50 LINCS-CP signatures most negatively correlated with the LAM signature (columns) with the corresponding matched genes (rows) are presented via heatmap. **B)** Odds ratios of the top most enriched MOA categories are shown via dot plot where the size of the dots represents the significance of the MOA categories with a bigger dot indicating lower FDR adjusted P-value. MOA categories were selected based on odds ratio>1.75, −log10(FDR)>4, and at least 150 signatures in any MOA category. **C)** Distribution of the overall signature correlations associated with each of the MOA categories are demonstrated via box-and-whisker plots. Each dot represents a LINCS-CP signature and negative correlations indicate the potential of the drug mechanisms to revert the LAM signature.

## Discussion

The connectivity analysis leveraging large databases of transcriptional perturbation signatures such as LINCS-L1000 along with the open accessibility to processed transcriptomics data^43,44^ and signatures^45,46^, enables *in silico* discovery of novel therapeutics. However, disease-related biological processes and resulting transcriptional dysregulation are not uniform across all cell types within the diseased tissues. Furthermore, the differences in expression profiles between cells of different types usually dwarf the differences between diseased and non-diseased cells of the same type. Therefore, the cell-averaging in the traditional bulk assays can produce disease transcriptional signatures of no relevance for finding putative therapeutics via connectivity analysis. This has been clearly demonstrated in our analysis of LAM data.

The scRNA-seq data used in our analysis was previously described and analyzed by Guo *et al.*^20^, and our pathway analysis results of naïve LAM signatures are consistent with results presented in that paper. Unlike Guo *et al.*, we were also able to identify a small set of cells expressing known LAM markers in the sirolimus treated LAM sample. However, the most important contribution of our study is the connectivity analysis of the LAM signatures.

Identification of remission-inducing therapeutic agents that can eliminate LAM cells has been challenging. Our connectivity analyses identified several known and novel repurposable therapeutic agents for LAM treatment including mTORC1 inhibitors as potential therapeutics to revert the LAM signature. However, mTORC1 inhibitors were not enriched among the connected MOAs indicating that the bulk transcriptional signatures cannot capture the key driving molecular mechanism of LAM. This was further supported by the pathway enrichments where genes up-regulated through activation of mTORC1 complex were enriched in the single cell DCS of naïve LAM, but not in the bulk signature. This demonstrates the importance of single cell profiling and effectiveness of our proposed workflow for scRNA-seq based connectivity analysis. To the best of our knowledge, this is the first analysis that describes and clearly demonstrates the importance of single cell transcriptional signature based connectivity analysis.

In addition to mTORC1 inhibitors, our analysis also identified additional classes of drugs, as well as specific drugs, capable of reverting the LAM signature such as, antiproliferative CDK inhibitors, and MEK/MAPK inhibitors, which might induce cytotoxicity against the LAM cells. The analysis of sirolimus treated LAM, implicated NF-κB pathway and proteasome inhibitors which have already been considered as therapeutic strategy in TSC. Functional enrichments of sirolimus treated LAM signature identified interferon gamma response which might lead to the activation of pro-survival pathways such as NF-κB. Furthermore, other cellular processes such as response to oxidative stress and antigen processing and presentation were induced in LAM2 signature implicating strong connectivity of NF-κB pathway and proteasome inhibitors. Additionally, several ongoing trials are testing the efficacy of multi-kinase inhibitor, Src inhibitor, and Cyclooxygenase inhibitors in LAM have also been strongly implicated in our connectivity analysis confirming again relevancy of the analysis results.

## Methods

### Single-cell RNA-seq and LINCS-L1000 data

Single-cell RNA-seq (scRNA-seq) was performed on dissociated lung tissue samples that were collected from three different sources including an untreated LAM patient (LAM1), patient treated with sirolimus (LAM2), and a brain dead, beating-heart, organ donor control patient (WT). Both LAM patients were undergoing lung transplantation. Single-cell suspensions of the two explanted LAM lungs and the normal lung were subjected to 10x Chromium scRNA-seq. CellRanger pipeline was used for read alignment and quantification. Raw gene counts data used in this analysis have been previously described and submitted to GEO^20^ (GSE135851). LAM1 data corresponds to the sample GSM4035465, LAM2 data corresponds to sample GSM4035466 and WT sample corresponds to sample GSM4035472.

For connectivity analysis, we utilized LINCS-L1000 database which is comprised of an extensive library of over a million gene expression profiles^19^. L1000 assay, a low-cost high-throughput technology developed by the Broad Institute, measures the expression of 978 landmark genes. The gene expression profiles were generated in response to a wide range of perturbing agents including ~20,000 small molecule compounds in more than 100 human cell lines and cell types for a total of 473,647 signatures^18^. We considered 143,374 chemical perturbation signatures available via iLINCS^45^ which were constructed by merging level-4 L1000 signature replicates into level-5 moderated Z-scores and only the reproducible signatures were retained.

### Single-cell RNA-seq data pre-processing and clustering

For scRNA-seq data, we filtered low-quality cells that were expressed (unique molecular identifies (UMI)>0) in less than 500 genes and had more than 10% mitochondrial UMI counts. Initial data preprocessing, normalization, and clustering was performed using Seurat3^21^ for LAM1, LAM2, and WT samples individually. Data were normalized by the global-scaling normalization method (“LogNormalize”) and top 2000 genes with highest standardized variance (method=“vst”) were selected for principle component (PC) analysis. For clustering, shared nearest-neighbor (SNN) graph was constructed with top 30 PCs with highest variances and Louvain algorithm for community detection^47^ with resolution parameter of 0.8 was used for clustering of cells within each sample. For integrated clustering of LAM1 and LAM2, both samples were merged using “IntegrateData” based on the anchors from “FindIntegrationAnchors” object with default parameters in Seurat3. Resolution parameter was set to 0.4 for cell clustering in the integrated LAM.

### Construction of cluster annotating and disease characterizing signatures

We employed a two-step strategy to annotate cell clusters and construct disease characterizing signature. In step 1, pairwise differential expression (DE) of each cluster was computed using MAST^48^ Bioconductor package which generated *n_t_* – 1 DE for each cluster (Supplementary Figure 1A), where *n_t_* is the number of clusters in sample *t.* For each pairwise comparison, we calculated π-score^49^ by multiplying log2 fold change (LFC) and negative logarithm of *P*-values (corrected for multiple testing using Benjamini-Hochberg (BH) method^50^). This can be written as:

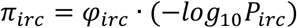

Where *φ_irc_* and *P_irc_* are LFC and *P*-values for *i^th^* gene, *r^th^* comparison, and *c^th^* cluster respectively. A positive *π* score indicates an up-regulation of a gene, whereas a negative score means down-regulation. A one-sided one sample Student’s *t*-test was carried out to combine the *n_t_* – 1 DEs into a cluster specific signature under the following hypotheses:

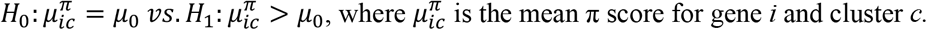

The null value was considered as 2 based on the cutoff of a gene being called differentially upregulated with pre-specified LFC of 1 and *P*-value of 0.01. *P*-values from *t-*test were further corrected for multiple testing using Benjamini-Hochberg method^50^. Top 200 most significantly (FDR<0.05) up-regulated genes were considered for cell-type/tissue enrichment via CLEAN^51^. The cluster of disease-critical LAM cells was identified as the one most enriched for 8 LAM marker genes.

In step 2, LAM specific cell cluster (LAMcluster16) was matched with WT clusters in terms of top 200 differentially upregulated (DU) genes (Supplementary Figure 1A). Similarities between LAM and WT clusters based on the number of overlapping genes were determined using complete linkage based hierarchical clustering with Euclidean distance measure. Significance of the overlaps among LAM and WT clusters were assessed via Fisher’s exact test. Finally, disease characterizing signature of both LAM1 and LAM2 were constructed by comparing LAM1 cells and LAM2 cells from LAM_cluster16_ with the matched WT clusters separately. Pseudo-bulk signatures for LAM1 and LAM2 were constructed by comparing all the LAM1 cells with WT cells and LAM2 cells with the WT cells respectively using MAST^48^ Bioconductor package.

### Connectivity analysis

LINCS-L1000 chemical perturbational (CP) signatures were considered for connectivity analysis. We selected 250 most significantly (FDR<0.05) differentially expressed (125 up-regulated and 125 down-regulated) genes from the LAM characterizing signature and matched them with the 978 L1000 landmark genes. Let, *Q_i_* be the LAM signature and *L_ij_* be the LINCS-CP signatures, where *i* is the set of matched genes and *j* is the set of LINCS CP signatures. Pearson correlation *Cor_j_*(*Q, L_j_*) was computed between LAM and each of the LINCS CP signatures (Supplementary Figure 1B) to assess the strength of relationship between the signatures. Negative correlation *P*-values were calculated for each signature correlation and corrected for multiple testing using BH method. A total of 86,538 LINCS CP signatures associated with 1005 unique mechanism of action (MOA) categories corresponding to the small molecules/drugs were considered for further MOA enrichment.

Let *M* be a binary variable where,

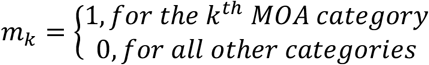

Here, *k* = 1,2,… …,1005. Inspired by the LRpath method^52^, we then fitted a small sample bias corrected binary logistic regression model^53^ for *M*,

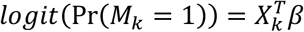

Where, negative logarithm of down-regulated *P*-values of correlation between LAM and LINCS-CP signatures is the predictor variable (Supplementary Figure 1B). *β* > 0 indicates that the signatures of the drugs for a specific MOA are “connected” with the disease signatures.

## Acknowledgements

This work was supported by the grants from National Institutes of Health: LINCS-BD2K DCIC (U54HL127624), Center for Environmental Genetics (P30ES006096) and the NHLBI research grant (R01HL138481); Department of Defense grant (W81XWH-19-1-0474) and by the Patient Benefit Grant Award from the LAM Foundation (LAM0133PB07-18).

## Author Contributions

N.A.M. developed the methodology and analyzed data, M.M. and J.Y. conceived the project, N.A.M. and M.M. conceived methodology, M.M. supervised methodology development and data analysis, E.Y.Z processed and validated tissues, S.S. assisted with single cell RNAseq, N.A.M, M.M. and J.Y. interpreted results and wrote the manuscript. All authors reviewed the manuscript.

