## Supplementary materials for "Connectivity analysis of single cell RNA-sequencing derived transcriptional signature of lymphangioleiomyomatosis"

### Supplementary Figures

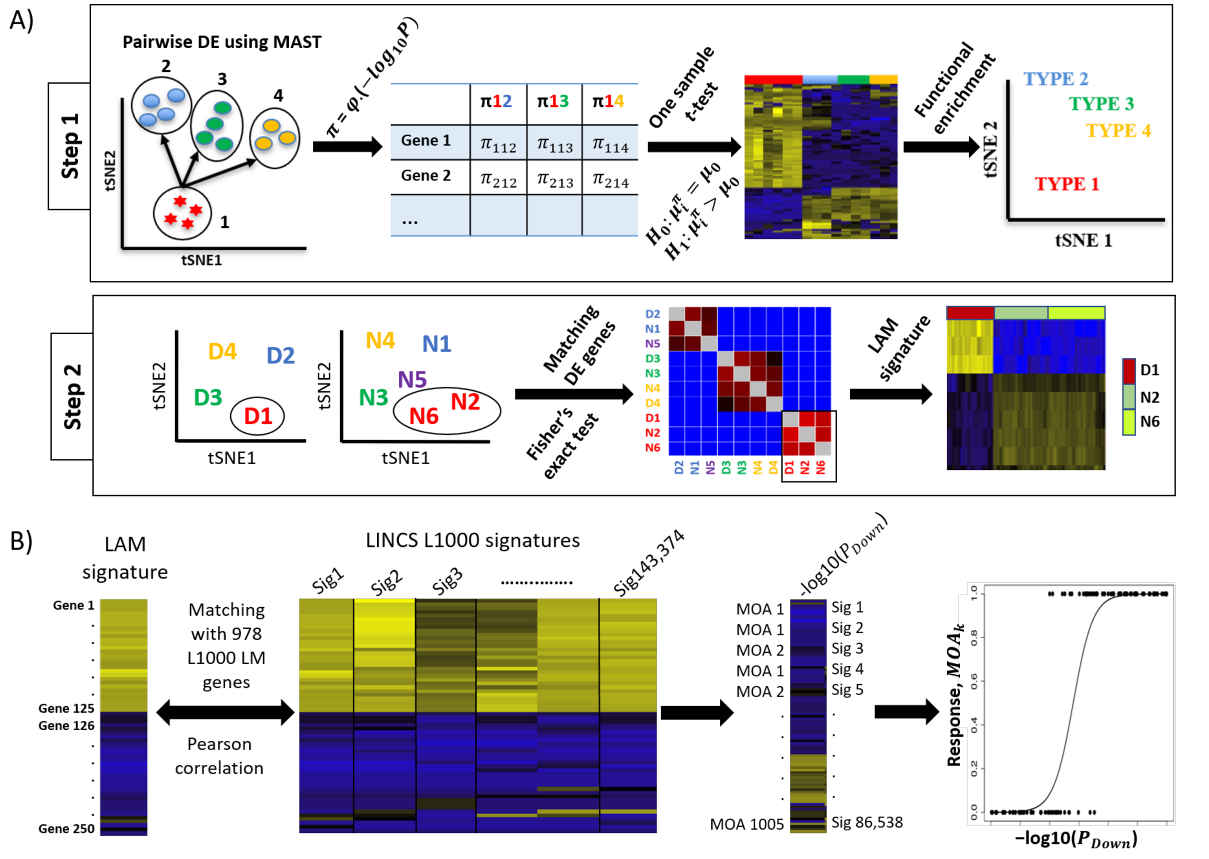


**Supplementary Figure 1. Conceptual framework for constructing LAM signature and connectivity analysis. A)** Hypothetical example illustrating identification of the cell types and signature construction of the LAM cell clusters. In step 1, pairwise differential expression (DE) is created for a single cluster (say cluster 1) such as, cluster 1 vs. cluster 2, cluster 1 vs. cluster 3 etc. using MAST Bioconductor package which generates a DE table with log2 fold changes ($\varphi$) and FDR-adjusted $P$-values (P) for each comparison and $\pi$ scores are calculated for each of the comparisons by multiplying $-log10(P)$ and $\varphi$. Pairwise DE results are then combined into a single DE by employing a one sample $t$-test of the $\pi$ scores for each gene under the hypotheses: $H_0: \mu_i^\pi = \mu_0$ vs. $H_1: \mu_i^\pi > \mu_0$, where null value is considered as 2 by considering P=0.01 and $\varphi$=1. Cell types are then identified by conducting functional enrichment of the top differentially upregulated (DU) genes. Step 1 is repeated for each of the clusters to identify corresponding cell types. In step 2, clusters from both LAM and normal samples are matched based on their DU genes and Fisher's exact test is used to test for cluster similarity. Finally, LAM cluster (say, D1) is then compared with the matching normal clusters (say, N1 and N6) to create the signature of the LAM cluster. **B)** For connectivity analysis, 125 most significantly up and 125 most significantly down regulated genes from LAM signature from **(A)** is matched with 978 LINCS-L1000 landmark genes and correlated with each of the 143,374 LINCS L1000 chemical perturbagen (CP) signatures using Pearson correlation. Down regulated FDR adjusted $P$-values (-log10 scale) of the correlations and corresponding mechanism of action (MOA) categories are extracted for LINCS-CP signatures. A total of 1005 unique MOAs are considered with the corresponding 86,538 signatures where each MOA has at least 1 associated signature. A binary logistic regression model with adjustment for small sample bias is then fitted for each of the MOA categories where negative logarithm of FDR-adjusted down-regulated $P$-values of the correlations between LAM and LINCS-CP signatures is considered as the predictor variable.

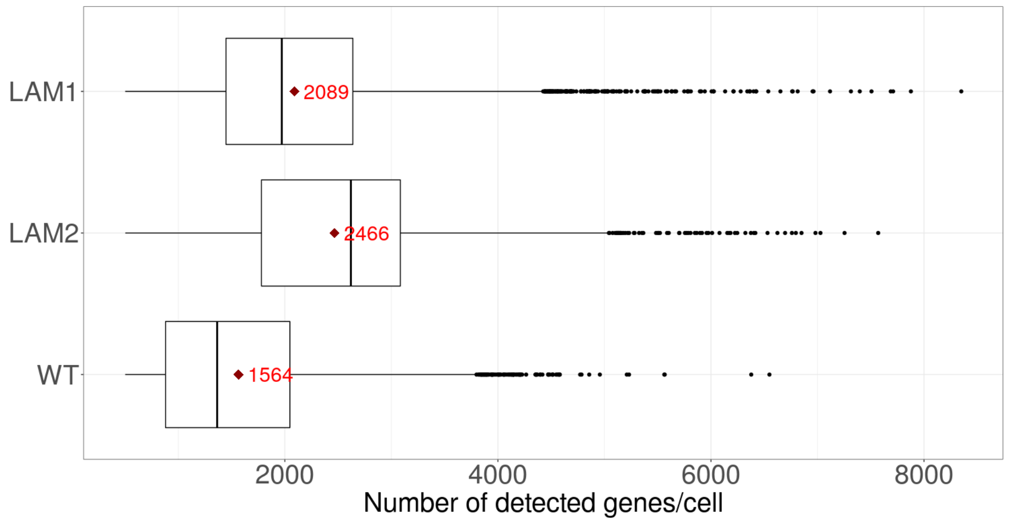


**Supplementary Figure 2. Distribution of the number of detected genes per cell.** Among the three samples, WT has the lowest average of detected genes per cell. Diamond sign indicates mean number of cells detected.

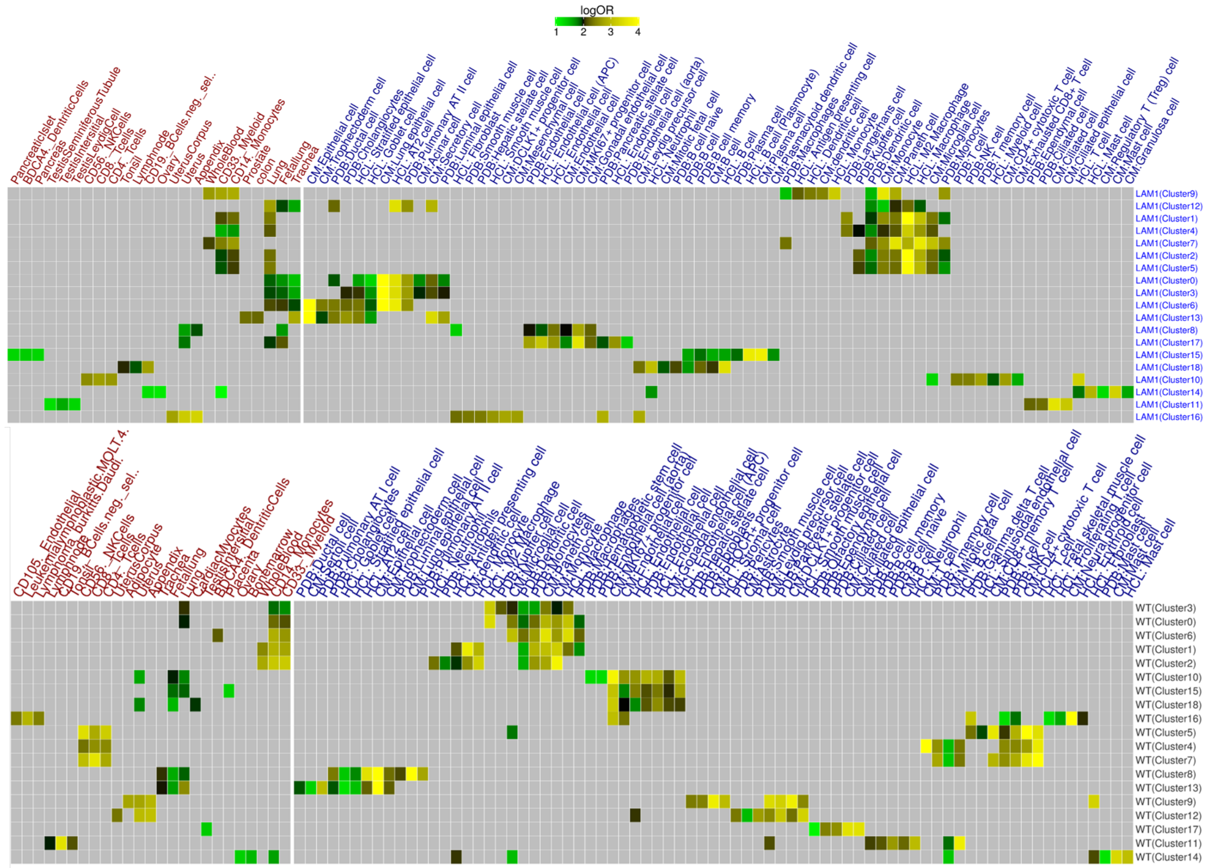


**Supplementary Figure 3. Hierarchical clustering of tissue and cell types enriched in LAM1 (top) and WT (bottom) clusters.** Top 200 most upregulated genes from each cluster were considered for tissue (colored in red) and cell type enrichment (colored in blue) and top three tissue categories from the gene atlas dataset and cell types from each of the three databases (CM, HCL, and PDB) were selected based on log OR.

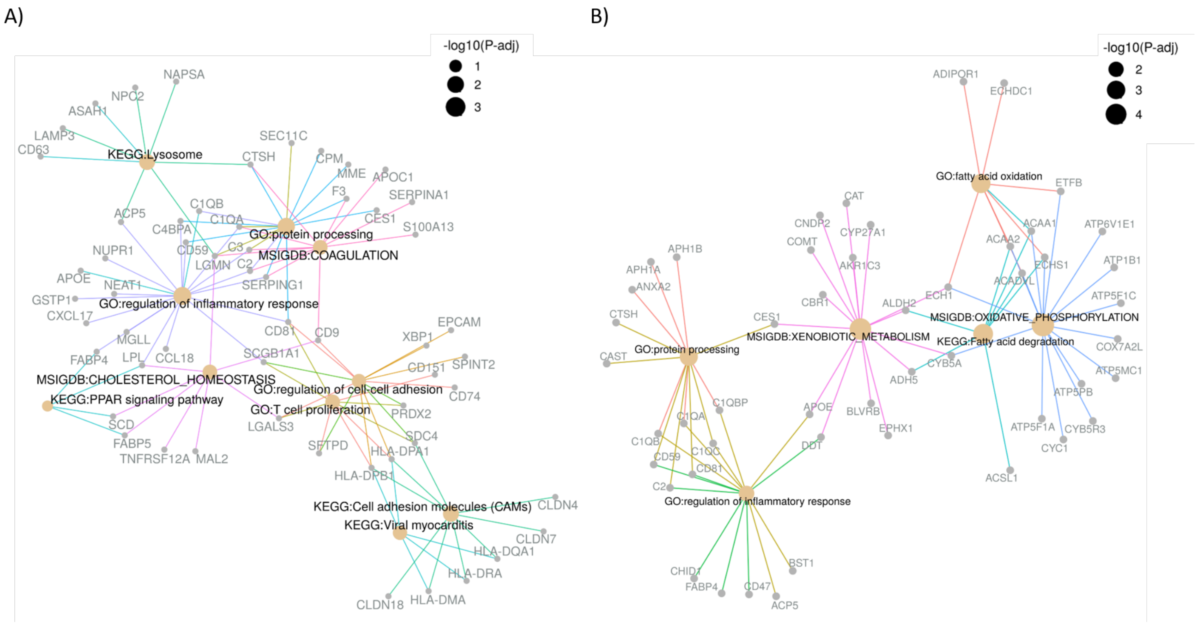


**Supplementary Figure 4. Functional enrichment of pseudo-bulk signatures.** Top 200 most significantly (FDR<0.05) up-regulated genes from bulk-LAM1 (**A**) and bulk-LAM2 (**B**) were considered for functional enrichment. Functional categories included GO (biological processes)/KEGG/MSigDB (Hallmark) and enrichment was conducted using clusterProfiler package. Enriched categories are represented by different edge colors and size of the nodes represent negative logarithm of FDR adjusted P-value.

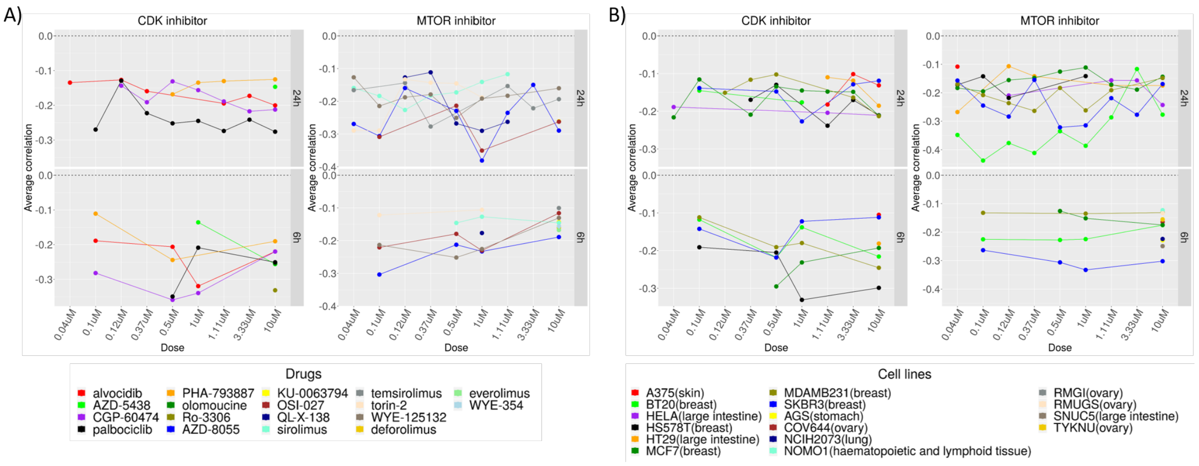


**Supplementary Figure 5. LAM1$_{cluster16}$ signature correlations at different levels of drug/cell-line/dose/time point.** Average correlations of top two MOA categories from LAM1$_{cluster16}$ are shown via line plots at different levels of dosages (in μM) and time points (in hours) with respect to drugs (**A**) and cell lines (**B**). Each dot represents a LINCS-CP signature and it's correlation with the LAM signature where drugs/cell-lines are coded by distinct colors. Signatures were further filtered with correlation below -0.1 and only female cell lines were considered where primary tissue sites are indicated within parenthesis in (**B**).

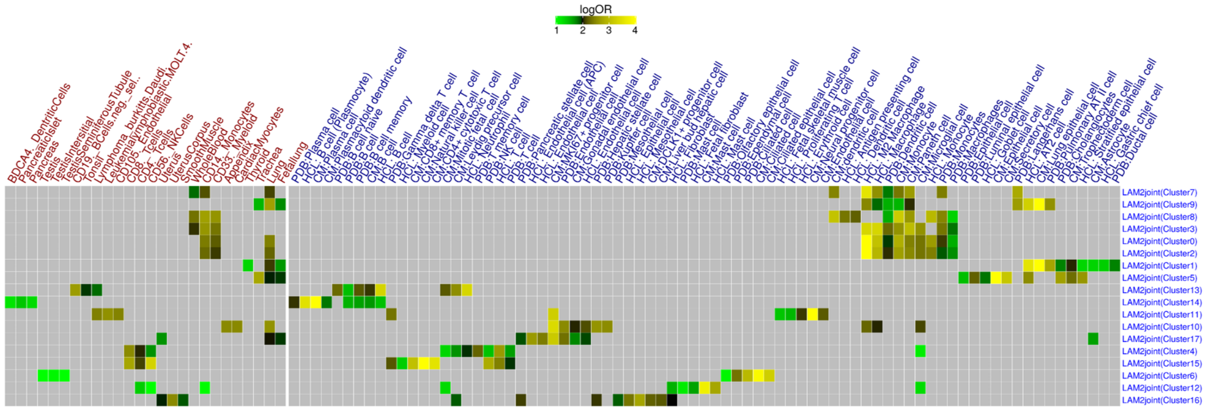


**Supplementary Figure 6. Hierarchical clustering of tissue and cell types enriched in joint-LAM2 clusters.** Top 200 most upregulated genes from each cluster were considered for tissue (colored in red) and cell type enrichment (colored in blue) and top three tissue categories from the gene atlas dataset and cell types from each of the three databases (CM, HCL, and PDB) were selected based on log OR.

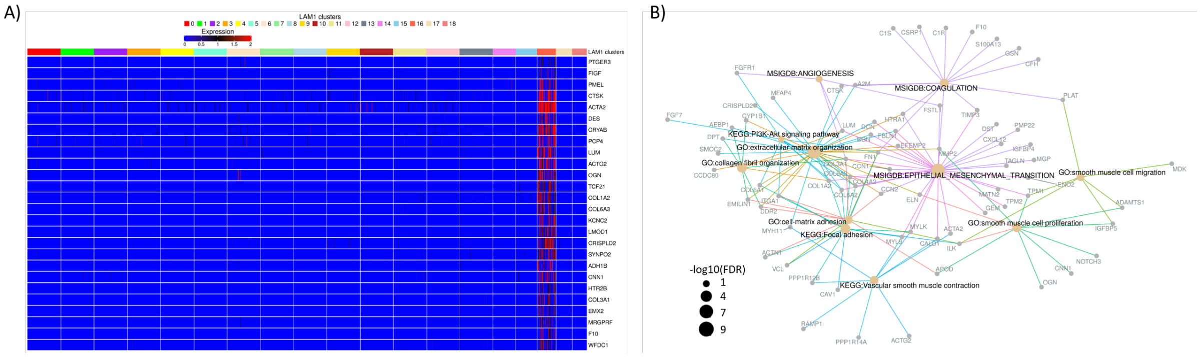


**Supplementary Figure 7. Selected differentially and distinctively expressed genes in LAM cell clusters and corresponding functional enrichment. A)** Top most differentially up-regulated (DU) and uniquely expressed genes in cluster annotating signature (CAS) of $LAM1_{cluster16}$. A gene is considered DU with an FDR adjusted P-value<0.05 and t-statistic value>0. Clusters were down-sampled to 100 using Seurat's "subset" function. Top 200 DU genes were further subsetted with an average expression above $30^{th}$ percentile in $LAM1_{cluster16}$ and below $10^{th}$ percentile in all other clusters were considered as differentially and distinctively expressed (DDE). **B)** Top 200 most significantly (FDR<0.05) up-regulated genes from the cluster annotating signature of $LAM1_{cluster16}$ were considered for functional enrichment. Functional categories included GO (biological processes)/KEGG/MSigDB (Hallmark) and enrichment was conducted using clusterProfiler package. Enriched categories are represented by different edge colors and size of the nodes represent negative logarithm of FDR adjusted P-value.

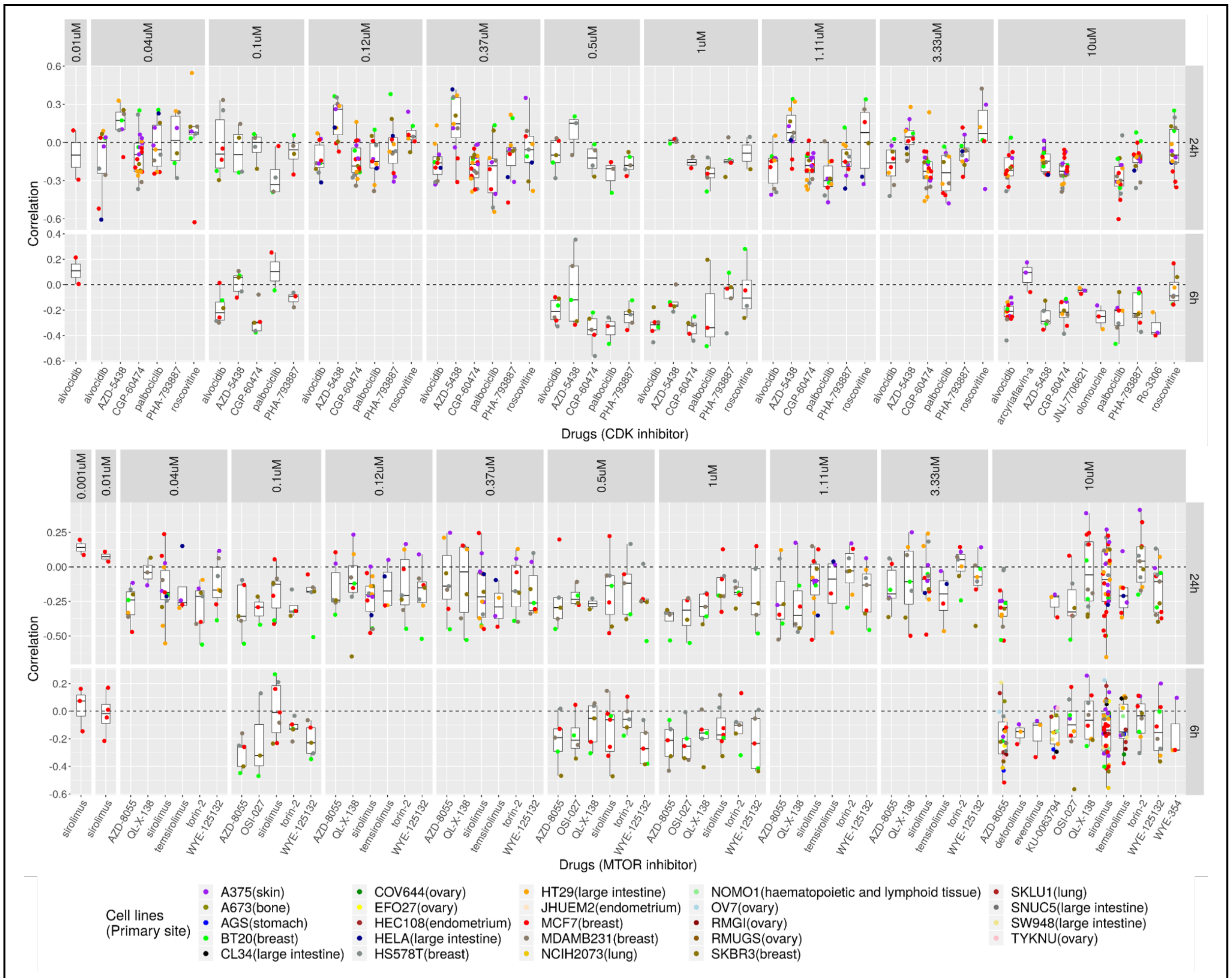


**Supplementary Figure 8. Disease characterizing signature correlations of LAM1 at individual drugs/cell lines/dosages/time points.** Correlation distribution of top 2 MOA categories are presented in the box-and-whisker plots at different levels of drugs, cell lines, concentrations, and time points. Cell lines with corresponding primary tissue sites are indicated by distinct colors. Bottom, left, top, and right axes represent drugs/small molecules, correlation, dosage concentrations (in µM), and time points (in hours) respectively. Each dot represents a LINCS-CP signature and it's correlation with the LAM1 signature.

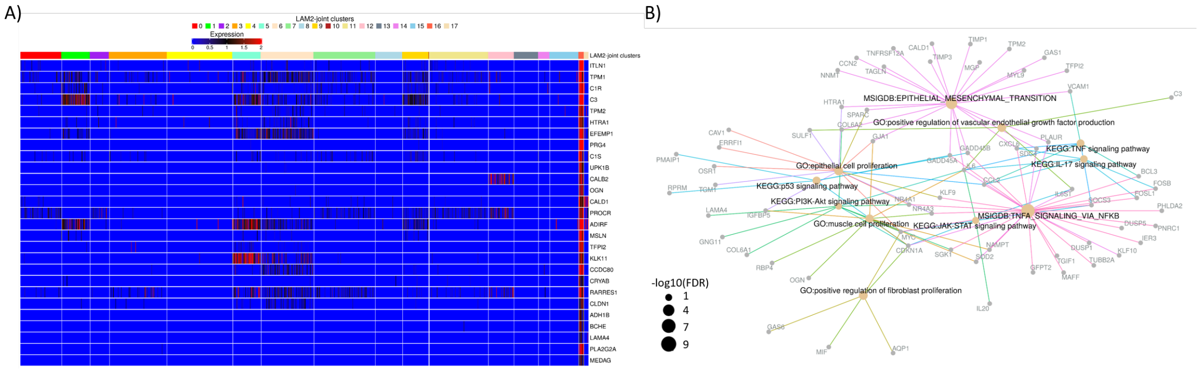


**Supplementary Figure 9. Selected differentially and distinctively expressed genes in LAM2$_{\text{joint-cluster16}}$ cell clusters and corresponding functional enrichment. A)** Top most differentially up-regulated (DU) and uniquely expressed genes in cluster annotating signature (CAS) of LAM2$_{\text{joint-cluster16}}$. A gene is considered DU with an FDR adjusted P-value<0.05 and t-statistic value>0. Clusters were down-sampled to 100 using Seurat's "subset" function. Top 200 DU genes were further subsetted with an average expression above 30$^{\text{th}}$ percentile in LAM2$_{\text{joint-cluster16}}$ and below 10$^{\text{th}}$ percentile in all other clusters were considered as differentially and distinctively expressed (DDE). **B)** Top 200 most significantly (FDR<0.05) up-regulated genes from the cluster annotating signature of LAM2$_{\text{joint-cluster16}}$ were considered for functional enrichment. Functional categories included GO (biological processes)/KEGG/MSigDB (Hallmark) and enrichment was conducted using clusterProfiler package. Enriched categories are represented by different edge colors and size of the nodes represent negative logarithm of FDR adjusted P-value.

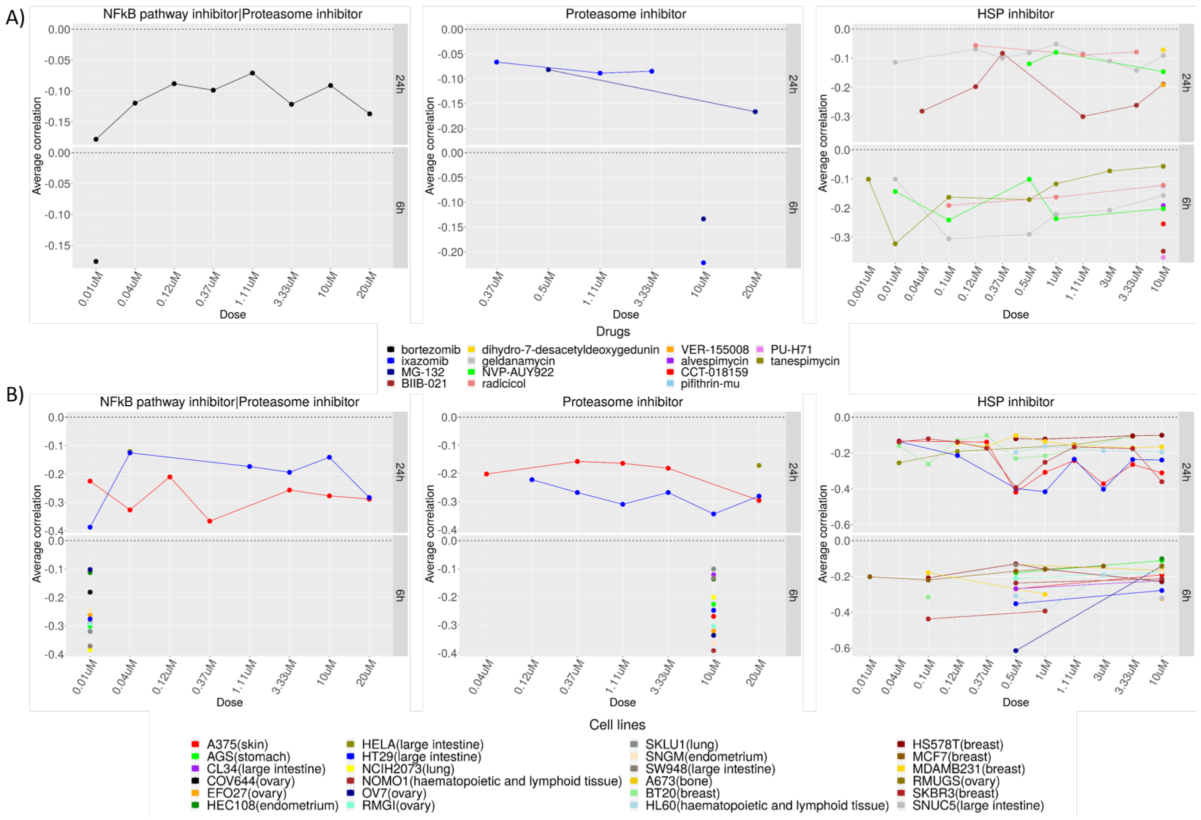


**Supplementary Figure 10. LAM2 signature correlations at different levels of drug/cell-line/dose/time point.** Average correlations of top three MOA categories from $LAM2_{joint\text{-}cluster16}$ are shown via line plots at different levels of dosages (in μM) and time points (in hours) with respect to drugs (**A**) and cell lines (**B**). Each dot represents a LINCS-CP signature and it's correlation with the LAM signature where drugs/cell-lines are coded by distinct colors. Signatures were further filtered with correlation below -0.05 and only female cell lines were considered where primary tissue sites are indicated within parenthesis in (**B**).

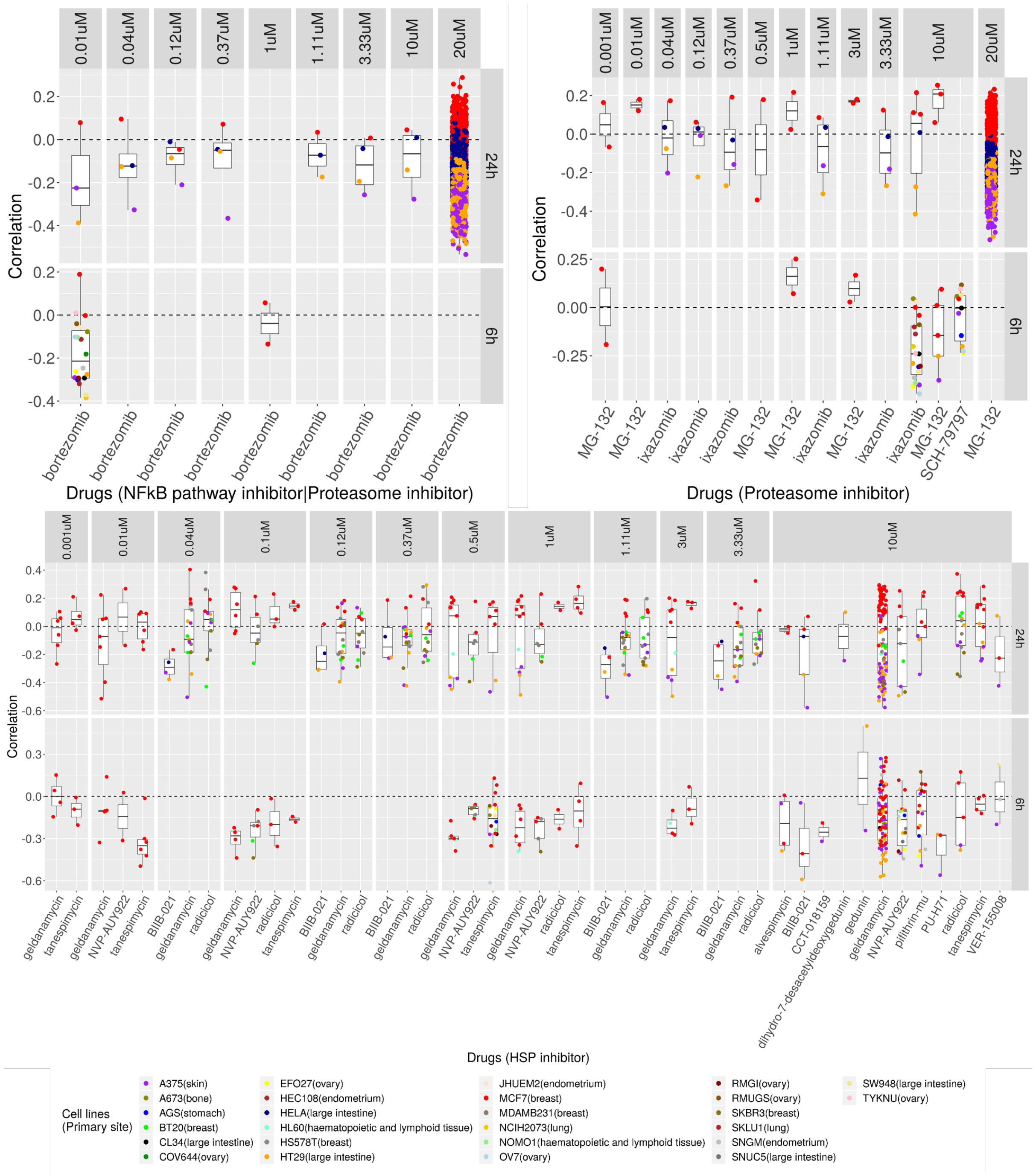


**Supplementary Figure 11. Disease characterizing signature correlations of LAM2 at individual drugs/cell lines/dosages/time points. A)** Correlation distribution of top 3 MOA categories are presented in the box-and-whisker plots at different levels of drugs, cell lines, concentrations, and time points. Cell lines with corresponding primary tissue sites are indicated by distinct colors. Bottom, left, top, and right axes represent drugs/small molecules, correlation, dosage concentrations (in µM), and time points (in hours) respectively. Each dot represents a LINCS-CP signature and it's correlation with the LAM2 signature.

### Supplementary tables

| Markers | Reference |
|---|---|
| PTGER3 | Li et al., 2017 [1] |
| PMEL | Hoon et al., 1994[2] |
| PCP4 | Guo et al., 2020[3] |
| CTSK | Chilosi et al., 2009[4] |
| ACTA2 | Zhe et al., 2004[5], Matsui et al., 2001[6] |
| FIGF | Seyama et al., 2006[7] |
| DES | Matsui et al., 2001[6], Taveira-DaSilva et al., 2006[8] |
| CRYAB | Iwaki and Tateishi, 1991[9] |

**Supplementary Table 1. List of known LAM markers.** A set of 8 markers associated with LAM are curated from the literature.

| | OR LAM1 | PValue LAM1 | OR LAM2 | PValue LAM2 | OR JointLAM | PValue JointLAM | OR JointLAM2 | PValue JointLAM2 |
|---|---|---|---|---|---|---|---|---|
| Cluster:0 | 0 | 1 | 0 | 1 | 0 | 1 | 0 | 1 |
| Cluster:1 | 0 | 1 | 0 | 1 | 0 | 1 | 0 | 1 |
| Cluster:2 | 0 | 1 | 0 | 1 | 0 | 1 | 0 | 1 |
| Cluster:3 | 0 | 1 | 0 | 1 | 2.1227 | 0.4063 | 3.4012 | 0.28075 |
| Cluster:4 | 0 | 1 | 0 | 1 | 0 | 1 | 0 | 1 |
| Cluster:5 | 0 | 1 | 0 | 1 | 0 | 1 | 0 | 1 |
| Cluster:6 | 0 | 1 | 2.7848 | 0.3301 | 0 | 1 | 0 | 1 |
| Cluster:7 | 0 | 1 | 0 | 1 | 0 | 1 | 0 | 1 |
| Cluster:8 | 0 | 1 | 0 | 1 | 0 | 1 | 0 | 1 |
| Cluster:9 | 0 | 1 | 0 | 1 | 0 | 1 | 0 | 1 |
| Cluster:10 | 0 | 1 | 0 | 1 | 0 | 1 | 0 | 1 |
| Cluster:11 | 0 | 1 | 0 | 1 | 0 | 1 | 0 | 1 |
| Cluster:12 | 0 | 1 | 2.2081 | 0.3947 | 0 | 1 | 0 | 1 |
| Cluster:13 | 0 | 1 | 0 | 1 | 0 | 1 | 0 | 1 |
| Cluster:14 | 0 | 1 | 0 | 1 | 0 | 1 | 0 | 1 |
| Cluster:15 | 0 | 1 | 0 | 1 | 0 | 1 | 0 | 1 |
| Cluster:16 | Inf | 1.32E-09 | 0 | 1 | Inf | 3.31E-10 | 48.538 | 2.14E-06 |
| Cluster:17 | 0 | 1 | 0 | 1 | 0 | 1 | 0 | 1 |
| Cluster:18 | 0 | 1 | 0 | 1 | NA | NA | NA | NA |

**Supplementary Table 2. Enrichment of 8 LAM markers in LAM1, LAM2-joint, and LAM-integrated clusters.** Significantly differentially up-regulated genes (FDR<0.05) were considered for marker enrichment. Odds ratio (OR) and P-values were calculated using Fisher's exact test. Infinite OR means all the 8 markers are present in the signature of the corresponding cluster.

| MOA | Estimate | PValue | FDR | OR | 95% lower | 95% upper | No. of signatures | No. of compounds |
|---|---|---|---|---|---|---|---|---|
| CDK inhibitor | 0.863 | 2.89E-85 | 4.31E-83 | 2.371 | 2.174 | 2.585 | 1701 | 15 |
| MTOR inhibitor | 0.83 | 5.85E-75 | 4.36E-73 | 2.314 | 2.115 | 2.531 | 1592 | 12 |
| PI3K inhibitor | 0.630 | 2.65E-65 | 1.32E-63 | 1.878 | 1.747 | 2.019 | 2922 | 19 |
| HSP inhibitor | 0.634 | 3.06E-45 | 1.14E-43 | 1.885 | 1.727 | 2.059 | 1904 | 14 |
| PI3K inhibitor\|MTOR inhibitor | 0.974 | 4.04E-33 | 1.2E-31 | 2.649 | 2.259 | 3.106 | 428 | 2 |
| HDAC inhibitor | 0.461 | 7.13E-33 | 1.77E-31 | 1.587 | 1.471 | 1.712 | 2947 | 41 |
| MEK inhibitor\|MAP kinase inhibitor\|Protein kinase inhibitor | 0.703 | 7.97E-22 | 1.7E-20 | 2.02 | 1.75 | 2.332 | 652 | 1 |
| MTOR inhibitor\|DNA dependent protein kinase inhibitor\|Phosphodiesterase inhibitor\|PLK inhibitor | 1.004 | 1.61E-21 | 3E-20 | 2.73 | 2.22 | 3.357 | 244 | 1 |
| RNA polymerase inhibitor | 1.022 | 2.74E-15 | 4.54E-14 | 2.78 | 2.157 | 3.582 | 158 | 6 |
| CDK inhibitor\|Cell cycle inhibitor\|MCL1 inhibitor | 0.947 | 3.34E-13 | 4.52E-12 | 2.579 | 1.999 | 3.329 | 166 | 1 |
| RAF inhibitor | 0.450 | 2.49E-12 | 3.09E-11 | 1.57 | 1.384 | 1.781 | 1021 | 9 |
| Glucocorticoid receptor agonist | 0.422 | 1.83E-10 | 2.09E-09 | 1.526 | 1.34 | 1.738 | 980 | 42 |
| HDAC inhibitor\|CDK expression enhancer\|ID1 expression inhibitor | 0.384 | 2.73E-08 | 2.14E-07 | 1.469 | 1.283 | 1.682 | 924 | 1 |
| AKT inhibitor | 0.417 | 3.89E-08 | 2.9E-07 | 1.519 | 1.308 | 1.763 | 744 | 8 |
| EGFR inhibitor\|JAK inhibitor\|Leukotriene inhibitor\|Mediator release inhibitor | 0.82 | 8.47E-08 | 6.01E-07 | 2.273 | 1.683 | 3.069 | 132 | 1 |
| MAP kinase inhibitor\|Protein kinase inhibitor | 0.885 | 1.34E-07 | 9.07E-07 | 2.424 | 1.744 | 3.369 | 104 | 1 |
| Aurora kinase inhibitor\|Bcr-Abl kinase inhibitor\|FLT3 inhibitor\|JAK inhibitor | 0.454 | 9.13E-05 | 0.00041 | 1.575 | 1.255 | 1.978 | 307 | 1 |
| RNA synthesis inhibitor\|Topoisomerase inhibitor | 0.610 | 9.35E-05 | 0.00041 | 1.841 | 1.355 | 2.5 | 150 | 1 |
| VEGFR inhibitor | 0.346 | 0.000198 | 0.00082 | 1.414 | 1.178 | 1.698 | 518 | 9 |
| CLK2 inihibitor | 0.589 | 0.000269 | 0.00104 | 1.803 | 1.313 | 2.475 | 142 | 1 |

| | | | | | | | | |
|---|---|---|---|---|---|---|---|---|
| EPHB3 inhibitor | 0.659 | 0.000272 | 0.00104 | 1.934 | 1.356 | 2.757 | 107 | 1 |
| ITK inhibitor | 0.604 | 0.000878 | 0.002845 | 1.83 | 1.282 | 2.613 | 111 | 2 |
| FLT3 inhibitor\|KIT inhibitor\|PDGFR tyrosine kinase receptor inhibitor\|RAD51 inhibitor\|RET tyrosine kinase inhibitor | 0.515 | 0.001263 | 0.00384 | 1.674 | 1.224 | 2.289 | 154 | 1 |
| CDK inhibitor\|Cell cycle inhibitor | 0.425 | 0.001835 | 0.004965 | 1.53 | 1.171 | 2 | 226 | 1 |
| IGF-1 inhibitor | 0.356 | 0.00302 | 0.007143 | 1.428 | 1.129 | 1.808 | 307 | 6 |
| Topoisomerase inhibitor | 0.187 | 0.003463 | 0.007938 | 1.207 | 1.064 | 1.368 | 1229 | 22 |
| Inositol monophosphatase inhibitor | 0.496 | 0.005217 | 0.011265 | 1.643 | 1.16 | 2.327 | 126 | 3 |
| Calcineurin inhibitor | 0.403 | 0.00676 | 0.013799 | 1.497 | 1.118 | 2.003 | 193 | 3 |
| ALK tyrosine kinase receptor inhibitor | 0.235 | 0.033956 | 0.043968 | 1.266 | 1.018 | 1.574 | 391 | 2 |

**Supplementary Table 3. MOA enrichment of disease characterizing signature of LAM1$_{\text{cluster16}}$.** Categories are selected with at least 100 signatures.

| MOA | Estimate | PValue | FDR | OR | 95% lower | 95% upper | No. of signatures | No. of compounds |
|---|---|---|---|---|---|---|---|---|
| NFkB pathway inhibitor\|Proteasome inhibitor | 1.389 | 0 | 0 | 4.011 | 3.741 | 4.3 | 2214 | 1 |
| Proteasome inhibitor | 1.466 | 0 | 0 | 4.333 | 4.049 | 4.636 | 2345 | 5 |
| HSP inhibitor | 0.823 | 6.09E-85 | 3.207E-83 | 2.276 | 2.096 | 2.472 | 1904 | 14 |
| MTOR inhibitor | 0.596 | 7.04E-34 | 2.783E-32 | 1.814 | 1.648 | 1.998 | 1592 | 12 |
| Farnesyltransferase inhibitor\|NFkB pathway inhibitor | 1.267 | 2E-24 | 6.311E-23 | 3.552 | 2.784 | 4.531 | 136 | 1 |
| KIT inhibitor\|Src inhibitor\|Bcr-Abl kinase inhibitor\|Ephrin inhibitor\|PDGFR tyrosine kinase receptor inhibitor\|Tyrosine kinase inhibitor | 0.668 | 5.89E-20 | 1.55E-18 | 1.95 | 1.69 | 2.25 | 662 | 1 |
| PI3K inhibitor | 0.257 | 3.73E-10 | 5.89E-09 | 1.293 | 1.193 | 1.401 | 2922 | 19 |

| | | | | | | | | |
|---|---|---|---|---|---|---|---|---|
| Topoisomerase inhibitor | 0.356 | 3.56E-09 | 5.12E-08 | 1.427 | 1.268 | 1.606 | 1229 | 22 |
| EGFR inhibitor | 0.316 | 5.54E-09 | 7.298E-08 | 1.371 | 1.233 | 1.525 | 1575 | 30 |
| PI3K inhibitor\|MTOR inhibitor | 0.54 | 1.24E-08 | 1.513E-07 | 1.715 | 1.425 | 2.065 | 428 | 2 |
| Cyclooxygenase inhibitor\|Histone acetyltransferase inhibitor\|Lipoxygenase inhibitor\|NFkB pathway inhibitor | 0.669 | 1.19E-06 | 1.174E-05 | 1.952 | 1.49 | 2.556 | 182 | 1 |
| RNA synthesis inhibitor\|Topoisomerase inhibitor | 0.688 | 4.74E-06 | 3.741E-05 | 1.989 | 1.482 | 2.671 | 150 | 1 |
| Glutathione transferase inhibitor | 0.719 | 3.8E-05 | 0.0002311 | 2.053 | 1.458 | 2.89 | 108 | 2 |
| AKT inhibitor | 0.316 | 5.26E-05 | 0.0002683 | 1.372 | 1.177 | 1.599 | 744 | 8 |
| Histone acetyltransferase inhibitor | 0.587 | 0.000728 | 0.0024098 | 1.799 | 1.28 | 2.529 | 121 | 2 |
| JAK inhibitor\|Lipocortin synthesis stimulant\|STAT inhibitor | 0.595 | 0.001448 | 0.0040124 | 1.812 | 1.257 | 2.613 | 104 | 1 |
| HDAC inhibitor\|CDK expression enhancer\|ID1 expression inhibitor | 0.225 | 0.001858 | 0.0049763 | 1.253 | 1.087 | 1.444 | 924 | 1 |
| CDK inhibitor\|PKC inhibitor | 0.545 | 0.002998 | 0.007288 | 1.724 | 1.203 | 2.47 | 112 | 1 |
| CDC inhibitor | 0.499 | 0.003509 | 0.00797 | 1.646 | 1.178 | 2.301 | 134 | 5 |
| PKC inhibitor | 0.283 | 0.003581 | 0.00797 | 1.327 | 1.097 | 1.606 | 489 | 9 |
| T-type calcium channel blocker | 0.514 | 0.005619 | 0.01153 | 1.672 | 1.162 | 2.405 | 112 | 3 |
| Retinoid receptor agonist | 0.264 | 0.006897 | 0.0135659 | 1.302 | 1.075 | 1.577 | 491 | 13 |
| Src inhibitor | 0.317 | 0.008101 | 0.0154219 | 1.373 | 1.086 | 1.737 | 313 | 8 |
| Protein synthesis inhibitor | 0.269 | 0.008716 | 0.0162515 | 1.309 | 1.07 | 1.6 | 443 | 19 |
| CLK2 inihibitor | 0.444 | 0.008743 | 0.0162515 | 1.559 | 1.119 | 2.173 | 142 | 1 |

| | | | | | | | | |
|---|---|---|---|---|---|---|---|---|
| RNA polymerase inhibitor | 0.408 | 0.012272 | 0.0217862 | 1.503 | 1.093 | 2.068 | 158 | 6 |
| Aurora kinase inhibitor\|Bcr-Abl kinase inhibitor\|FLT3 inhibitor\|JAK inhibitor | 0.294 | 0.015941 | 0.0262369 | 1.342 | 1.056 | 1.704 | 307 | 1 |
| DNA synthesis inhibitor | 0.343 | 0.028931 | 0.0426086 | 1.409 | 1.036 | 1.917 | 178 | 10 |
| NFkB pathway inhibitor | 0.212 | 0.032493 | 0.0454122 | 1.236 | 1.018 | 1.5 | 496 | 15 |
| Tubulin inhibitor | 0.182 | 0.035295 | 0.0475436 | 1.2 | 1.013 | 1.421 | 665 | 18 |
| Matrix metalloprotease inhibitor | 0.365 | 0.048328 | 0.0596547 | 1.441 | 1.003 | 2.07 | 126 | 5 |
| PPAR receptor agonist\|Insulin sensitizer | 0.24 | 0.062723 | 0.0734089 | 1.271 | 0.987 | 1.637 | 285 | 6 |
| PPAR receptor agonist | 0.254 | 0.063851 | 0.0736379 | 1.29 | 0.985 | 1.687 | 249 | 15 |
| Microtubule inhibitor\|Tubulin inhibitor | 0.327 | 0.069104 | 0.0773955 | 1.387 | 0.975 | 1.973 | 137 | 2 |
| Nucleophosmin inhibitor | 0.318 | 0.069558 | 0.0773955 | 1.375 | 0.975 | 1.939 | 145 | 7 |
| RAF inhibitor\|FLT3 inhibitor\|KIT inhibitor\|PDGFR tyrosine kinase receptor inhibitor\|RET tyrosine kinase inhibitor\|VEGFR inhibitor | 0.268 | 0.071671 | 0.0791888 | 1.308 | 0.977 | 1.751 | 209 | 1 |
| HMGCR inhibitor | 0.216 | 0.073729 | 0.0808975 | 1.241 | 0.979 | 1.573 | 330 | 11 |
| MTOR inhibitor\|DNA dependent protein kinase inhibitor\|Phosphodiesterase inhibitor\|PLK inhibitor | 0.235 | 0.092521 | 0.0935514 | 1.265 | 0.962 | 1.662 | 244 | 1 |

**Supplementary Table 4. MOA enrichment of disease characterizing signature of LAM2$_{joint\text{-}cluster16}$.** Categories are selected with at least 100 signatures.

| MOA | Estimate | PValue | FDR | OR | 95% lower | 95% upper | No. of signatures | No. of compounds |
|---|---|---|---|---|---|---|---|---|
| HDAC inhibitor | 1.051 | 0 | 0 | 2.861 | 2.709 | 3.02 | 2947 | 41 |
| HDAC inhibitor\|CDK expression enhancer\|ID1 expression inhibitor | 1.606 | 0 | 0 | 4.981 | 4.582 | 5.415 | 924 | 1 |
| NFkB pathway inhibitor\|Proteasome inhibitor | 1.671 | 0 | 0 | 5.317 | 5.009 | 5.643 | 2214 | 1 |
| Proteasome inhibitor | 1.593 | 0 | 0 | 4.92 | 4.643 | 5.214 | 2345 | 5 |
| IKK inhibitor | 0.618 | 2.72E-32 | 4.67E-31 | 1.856 | 1.675 | 2.056 | 926 | 7 |
| Glutathione transferase inhibitor | 1.318 | 5.29E-29 | 7.58E-28 | 3.736 | 2.965 | 4.708 | 108 | 2 |
| BTK inhibitor\|Cytoplasmic tyrosine protein kinase BMX inhibitor | 0.885 | 3.11E-17 | 3.34E-16 | 2.423 | 1.973 | 2.976 | 184 | 1 |
| Anti-inflammatory agent\|Antioxidant\|HSP inhibitor\|NFkB pathway inhibitor\|Topoisomerase inhibitor | 0.927 | 1.63E-16 | 1.55E-15 | 2.526 | 2.027 | 3.149 | 155 | 1 |
| Farnesyltransferase inhibitor\|NFkB pathway inhibitor | 0.964 | 3.54E-16 | 3.04E-15 | 2.623 | 2.08 | 3.307 | 136 | 1 |
| Topoisomerase inhibitor | 0.367 | 1.69E-13 | 1.32E-12 | 1.443 | 1.309 | 1.591 | 1229 | 22 |
| CLK2 inihibitor | 0.845 | 2.74E-12 | 1.96E-11 | 2.327 | 1.836 | 2.949 | 142 | 1 |
| JNK inhibitor | 0.41 | 1.27E-07 | 6.45E-07 | 1.506 | 1.294 | 1.753 | 482 | 6 |
| CDK inhibitor\|Cell cycle inhibitor\|MCL1 inhibitor | 0.597 | 1.11E-06 | 5.29E-06 | 1.816 | 1.429 | 2.309 | 166 | 1 |
| Mitochondrial DNA polymerase inhibitor\|Phosphatase inhibitor | 0.661 | 5.85E-06 | 2.33E-05 | 1.937 | 1.455 | 2.578 | 111 | 1 |
| CDK inhibitor | 0.204 | 5.96E-06 | 2.33E-05 | 1.226 | 1.122 | 1.339 | 1701 | 15 |
| NFkB pathway inhibitor | 0.35 | 7.61E-06 | 2.73E-05 | 1.419 | 1.217 | 1.653 | 496 | 15 |
| Nucleophosmin inhibitor | 0.582 | 9.8E-06 | 3.37E-05 | 1.79 | 1.383 | 2.317 | 145 | 7 |
| Histone acetyltransferase inhibitor | 0.59 | 4.03E-05 | 0.000133 | 1.803 | 1.361 | 2.389 | 121 | 2 |
| CDC inhibitor | 0.555 | 6.09E-05 | 0.000187 | 1.741 | 1.328 | 2.283 | 134 | 5 |
| KIT inhibitor\|Src inhibitor\|Bcr-Abl kinase inhibitor\|Ephrin inhibitor\|PDGFR tyrosine kinase receptor | 0.278 | 6.34E-05 | 0.000188 | 1.321 | 1.152 | 1.514 | 662 | 1 |

<table>
<tr><th></th><th></th><th></th><th></th><th></th><th></th><th></th><th></th><th></th></tr>
<tr><td>inhibitor|Tyrosine kinase inhibitor</td><td></td><td></td><td></td><td></td><td></td><td></td><td></td><td></td></tr>
<tr><td>Leucine rich repeat kinase inhibitor</td><td>0.467</td><td>8.11E-05</td><td>0.000232</td><td>1.595</td><td>1.264</td><td>2.011</td><td>196</td><td>1</td></tr>
<tr><td>HSP inhibitor</td><td>0.162</td><td>0.000169</td><td>0.000441</td><td>1.176</td><td>1.081</td><td>1.28</td><td>1904</td><td>14</td></tr>
<tr><td>Histone lysine methyltransferase inhibitor</td><td>0.514</td><td>0.000269</td><td>0.00068</td><td>1.671</td><td>1.268</td><td>2.203</td><td>133</td><td>5</td></tr>
<tr><td>JAK inhibitor|FLT3 inhibitor</td><td>0.402</td><td>0.00146</td><td>0.003393</td><td>1.494</td><td>1.167</td><td>1.914</td><td>181</td><td>1</td></tr>
<tr><td>T-type calcium channel blocker</td><td>0.464</td><td>0.003008</td><td>0.006633</td><td>1.59</td><td>1.171</td><td>2.161</td><td>112</td><td>3</td></tr>
<tr><td>Adenylyl cyclase activator</td><td>0.425</td><td>0.005147</td><td>0.010294</td><td>1.53</td><td>1.136</td><td>2.062</td><td>122</td><td>2</td></tr>
<tr><td>Cyclooxygenase inhibitor|Histone acetyltransferase inhibitor|Lipoxygenase inhibitor|NFkB pathway inhibitor</td><td>0.272</td><td>0.039503</td><td>0.054795</td><td>1.313</td><td>1.013</td><td>1.701</td><td>182</td><td>1</td></tr>
<tr><td>Leucine rich repeat kinase inhibitor|MAP kinase inhibitor</td><td>0.294</td><td>0.050196</td><td>0.065078</td><td>1.342</td><td>1</td><td>1.802</td><td>138</td><td>1</td></tr>
<tr><td>CHK inhibitor</td><td>0.179</td><td>0.080726</td><td>0.087879</td><td>1.196</td><td>0.978</td><td>1.461</td><td>327</td><td>3</td></tr>
<tr><td>Protein synthesis inhibitor</td><td>0.149</td><td>0.093682</td><td>0.094784</td><td>1.161</td><td>0.975</td><td>1.382</td><td>443</td><td>19</td></tr>
</table>

**Supplementary Table 5. MOA enrichment of disease characterizing pseudo-bulk signature of LAM1.** Categories are selected with at least 100 signatures.

<table>
<tr><th>MOA</th><th>Estimate</th><th>PValue</th><th>FDR</th><th>OR</th><th>95% lower</th><th>95% upper</th><th>No. of signatures</th><th>No. of compounds</th></tr>
<tr><td>NFkB pathway inhibitor|Proteasome inhibitor</td><td>0.588</td><td>0</td><td>0</td><td>1.801</td><td>1.775</td><td>1.828</td><td>2214</td><td>1</td></tr>
<tr><td>Proteasome inhibitor</td><td>0.55</td><td>0</td><td>0</td><td>1.733</td><td>1.709</td><td>1.758</td><td>2345</td><td>5</td></tr>
<tr><td>HDAC inhibitor|CDK expression enhancer|ID1 expression inhibitor</td><td>0.23</td><td>2.6E-94</td><td>6.68E-93</td><td>1.258</td><td>1.231</td><td>1.286</td><td>924</td><td>1</td></tr>
<tr><td>Protein synthesis inhibitor</td><td>0.282</td><td>8.89E-92</td><td>1.71E-90</td><td>1.326</td><td>1.29</td><td>1.362</td><td>443</td><td>19</td></tr>
<tr><td>ATPase inhibitor</td><td>0.252</td><td>1.37E-74</td><td>2.12E-73</td><td>1.287</td><td>1.253</td><td>1.322</td><td>515</td><td>26</td></tr>
<tr><td>IKK inhibitor</td><td>0.198</td><td>2.85E-61</td><td>3.14E-60</td><td>1.219</td><td>1.19</td><td>1.248</td><td>926</td><td>7</td></tr>
<tr><td>BTK inhibitor|Cytoplasmic tyrosine protein kinase BMX inhibitor</td><td>0.303</td><td>2E-52</td><td>1.72E-51</td><td>1.354</td><td>1.302</td><td>1.408</td><td>184</td><td>1</td></tr>
</table>

| | | | | | | | | |
|---|---|---|---|---|---|---|---|---|
| Farnesyltransferase inhibitor\|NFkB pathway inhibitor | 0.293 | 6.77E-36 | 4.74E-35 | 1.341 | 1.281 | 1.404 | 136 | 1 |
| HDAC inhibitor | 0.108 | 5.59E-35 | 3.59E-34 | 1.114 | 1.095 | 1.134 | 2947 | 41 |
| Glutathione transferase inhibitor | 0.307 | 9.97E-34 | 5.91E-33 | 1.36 | 1.294 | 1.429 | 108 | 2 |
| Mitochondrial DNA polymerase inhibitor\|Phosphatase inhibitor | 0.287 | 5.72E-28 | 2.94E-27 | 1.333 | 1.266 | 1.403 | 111 | 1 |
| Anti-inflammatory agent\|Antioxidant\|HSP inhibitor\|NFkB pathway inhibitor\|Topoisomerase inhibitor | 0.261 | 7.75E-28 | 3.73E-27 | 1.298 | 1.238 | 1.36 | 155 | 1 |
| PKC activator | 0.178 | 9.66E-15 | 3.54E-14 | 1.195 | 1.143 | 1.251 | 261 | 4 |
| JAK inhibitor\|Lipocortin synthesis stimulant\|STAT inhibitor | 0.234 | 3.49E-14 | 1.22E-13 | 1.264 | 1.19 | 1.343 | 104 | 1 |
| NFkB pathway inhibitor | 0.136 | 7.25E-13 | 2.43E-12 | 1.146 | 1.104 | 1.189 | 496 | 15 |
| CDC inhibitor | 0.187 | 1.95E-09 | 6.01E-09 | 1.206 | 1.134 | 1.282 | 134 | 5 |
| Histone lysine methyltransferase inhibitor | 0.185 | 3.61E-09 | 1.07E-08 | 1.204 | 1.132 | 1.28 | 133 | 5 |
| Histone acetyltransferase inhibitor | 0.149 | 4.71E-05 | 0.000104 | 1.16 | 1.08 | 1.247 | 121 | 2 |
| IKK inhibitor\|NFkB pathway inhibitor | 0.152 | 6.46E-05 | 0.000138 | 1.164 | 1.081 | 1.255 | 109 | 1 |
| T-type calcium channel blocker | 0.141 | 0.000273 | 0.000553 | 1.152 | 1.067 | 1.243 | 112 | 3 |
| HSP inhibitor | 0.043 | 0.000624 | 0.001202 | 1.044 | 1.019 | 1.071 | 1904 | 14 |
| Serine/threonine kinase inhibitor | 0.094 | 0.001467 | 0.002627 | 1.098 | 1.037 | 1.164 | 257 | 6 |
| Nucleophosmin inhibitor | 0.109 | 0.003621 | 0.006062 | 1.115 | 1.036 | 1.2 | 145 | 7 |
| Leucine rich repeat kinase inhibitor | 0.092 | 0.006877 | 0.011031 | 1.096 | 1.026 | 1.171 | 196 | 1 |
| Retinoid receptor agonist | 0.057 | 0.016584 | 0.026061 | 1.059 | 1.01 | 1.109 | 491 | 13 |
| LCK Inhibitor | 0.085 | 0.026646 | 0.040063 | 1.089 | 1.01 | 1.173 | 160 | 1 |
| CLK2 inihibitor | 0.089 | 0.027056 | 0.040063 | 1.093 | 1.01 | 1.182 | 142 | 1 |
| DNA synthesis inhibitor | 0.074 | 0.046431 | 0.063843 | 1.077 | 1.001 | 1.159 | 178 | 10 |
| Cyclooxygenase inhibitor\|Histone acetyltransferase inhibitor\|Lipoxygenase | 0.07 | 0.063003 | 0.079337 | 1.072 | 0.996 | 1.154 | 182 | 1 |

| | | | | | | | | |
|---|---|---|---|---|---|---|---|---|
| inhibitor\|NFkB pathway inhibitor | | | | | | | | |
| PLK inhibitor | 0.033 | 0.063718 | 0.079337 | 1.034 | 0.998 | 1.071 | 985 | 5 |
| DNA replication inhibitor\|STAT inhibitor | 0.077 | 0.066479 | 0.081252 | 1.08 | 0.995 | 1.173 | 138 | 1 |
| Adenosine receptor antagonist | 0.069 | 0.081253 | 0.089861 | 1.072 | 0.991 | 1.159 | 162 | 15 |

**Supplementary Table 6. MOA enrichment of disease characterizing pseudo-bulk signature of LAM2.** Categories are selected with at least 100 signatures.
